# NucBreak: Location of structural errors in a genome assembly by using paired-end Illumina reads

**DOI:** 10.1101/393488

**Authors:** Ksenia Khelik, Geir Kjetil Sandve, Alexander Johan Nederbragt, Torbjørn Rognes

## Abstract

**Background:** Advances in whole genome sequencing strategies have provided the opportunity for genomic and comparative genomic analysis of a vast variety of organisms. The analysis results are highly dependent on the quality of the genome assemblies used. Assessment of the assembly accuracy may significantly increase the reliability of the analysis results and is therefore of great importance.

**Results:** Here, we present a new tool called NucBreak aimed at detecting structural errors in assemblies, including insertions, deletions, duplications, inversions, and different inter-and intra-chromosomal rearrangements. NucBreak analyses the alignments of reads properly mapped to an assembly and exploits information about the alternative read alignments. We have compared NucBreak with other existing assembly accuracy assessment tools, namely Pilon, REAPR, and FRCbam as well as with several structural variant detection tools, including BreakDancer, Lumpy, and Wham, by using both simulated and real datasets.

**Conclusions:** The benchmarking results have shown that NucBreak in general predicts assembly errors of different types and sizes with relatively high sensitivity and with higher precision than the other tools. Such a balance between sensitivity and precision makes NucBreak a good alternative to the existing assembly accuracy assessment tools and SV detection tools. NucBreak is freely available at https://github.com/uio-bmi/NucBreak under the MPL license.

## 1. Background

Advances in whole genome sequencing technologies have led to a greatly increased number of organisms with sequenced genomes over the recent years. This has provided the opportunity to make genomic and comparative genomic analysis of a vast variety of organisms. The analysis results are highly dependent on the quality of the genome assemblies used. Any errors in an assembly directly impair analysis predictions and inferences based upon them [1]. The assessment of assembly accuracy may significantly increase the reliability of analysis results and is therefore of great importance.

There are several tools developed for genome assembly accuracy assessment, i.e. REAPR [2], FRCbam [3] and Pilon [4]. These tools identify regions with various inconsistencies in the alignments of reads mapped back to the assembly and detect the locations of assembly errors. The inconsistencies include abnormal read coverage, abnormal distance between reads in a pair relative to the insert size, wrong orientation of one or both reads in a pair, and a large percentage of soft-clipped reads (reads that are partly mapped to an assembly: the one end of the read is mapped to the reference while the second is not) and singletons (reads whose partner was not mapped). The tools are aimed at detecting structural errors including medium to long insertions and deletions, as well as inversions, duplications, and inter-and intra-chromosomal rearrangements. Pilon also enables detection of small insertions, deletions and substitutions and performs local assembly to fix detected assembly errors where possible.

The genome assembly accuracy assessment problem is very similar to the structural variant (SV) detection problem. The tools developed to detect structural variants between genomes of the same or closely related species, such as Wham [5], BreakDancer [6] and Lumpy [7], are based on the approaches similar to the ones implemented in REAPR, Pilon and FRCbam. They exploit the same types of inconsistencies in the read alignments in their workflow. The usage of such tools may be a possible alternative to the tools developed for genome assembly error detection.

Here we present a new tool NucBreak aimed at genome assembly accuracy assessment. In contrast to the other tools, it analyses the alignments of reads that are properly mapped to an assembly (where both reads in a pair are fully aligned in correct orientation at a reasonable distance) and exploits information about the alternative read alignments to detect the locations of assembly structural errors. The tool has been compared to REAPR, FRCbam and Pilon, the only existing tools detecting assembly error locations, as well as BreakDancer, Lumpy, and Wham. We have chosen BreakDancer, Lumpy, and Wham because they were developed to perform the analysis in whole genomes of different species and detect various types of structural variants compared to other existing SV detection tools. All tools have been tested for their ability to detect errors in assemblies by using either simulated or real datasets. The test results have shown that NucBreak enables prediction of assembly errors with higher precision than other tools, keeping relatively high level of sensitivity at the same time.

## 2. Implementation

NucBreak is a tool created to detect structural errors in an assembly by using paired-end Illumina reads. The reads are first mapped to the assembly, and then the mapping results are rigorously analysed to detect the assembly errors locations. The NucBreak workflow is shown in Figure 1.

**Figure 1:**
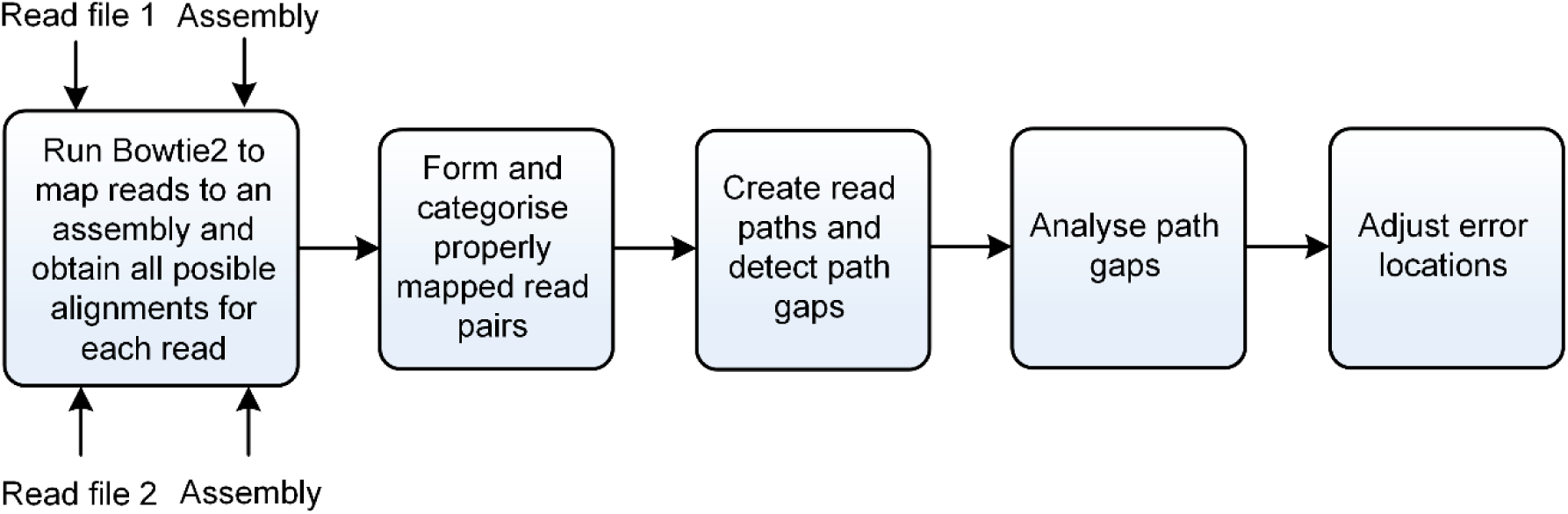
NucBreak workflow.

### 2.1 Read mapping

The error detection process starts with mapping reads to the assembly by using Bowtie2 [8]. Bowtie2 is run separately for each read file with the parameter settings “*--sensitive_local --ma 1 -a*” to report all local alignments with an added nucleotide match bonus. The resulting .sam files contain all possible alignments for each read not depending on the second read in a pair. A read alignment may contain either a full read sequence or a read sequence clipped on one or both ends. The read clipping is performed when one or both ends of a read accumulate a high number of differences compared to the assembly. The clipped part of a read may be mapped to another location in the assembly or remained unmapped. There may be a few short substitutions, insertions and/or deletions inside mapped reads or their parts.

### 2.2 Properly mapped read pair formation and categorization

Once the mapping results have been obtained, NucBreak combines reads into properly mapped read pairs and categorizes the pairs into several groups. A pair of reads is considered to be properly mapped if all of the five following conditions are fulfilled:

1. Both reads are mapped to the same assembly sequence.
2. The reads have different orientations relative to the assembly sequence.
3. The read with the reverse orientation is located at the same position or further down on the sequence compared to the mapping locations of the forward-oriented read.
4. The beginnings of the read sequences (the first bases of the read sequences as they are given in the input files) are not clipped. The exception is made only for the forward-oriented read mapped to the very beginning of the assembly sequence and the reverse-oriented read mapped to the very end of the assembly sequence.
5. The reads have a proper insert size (see [Additional file 1] for the details about the insert size detection approach).

The alignments of properly mapped reads may contain short substitutions, insertions and deletions.

To combine reads into properly mapped read pairs, NucBreak analyses all possible combinations of the read mapping locations for each input read pair and forms properly mapped read pairs from those reads whose locations satisfy the five conditions mentioned above. Each input read pair may give rise to none, one or several properly mapped read pairs (see Figure 2).

**Figure 2:**
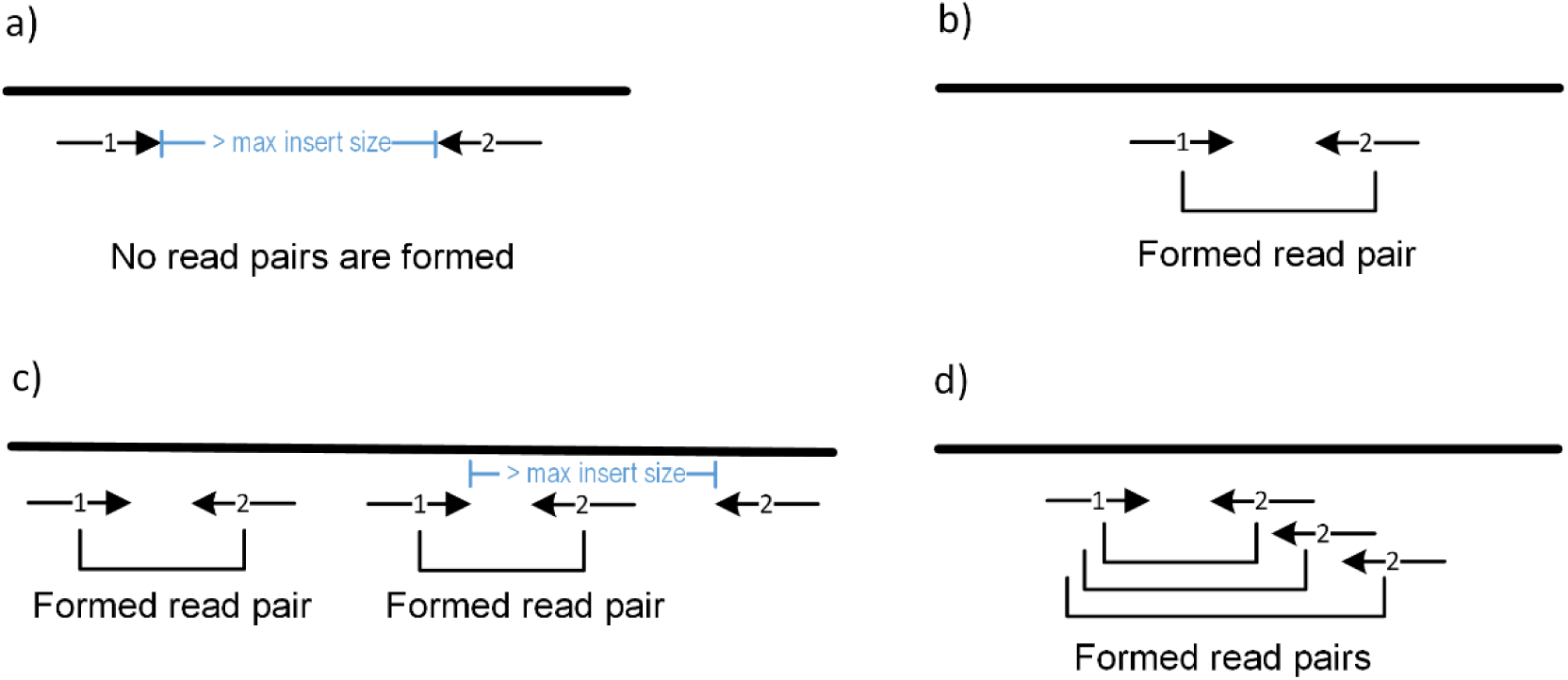
Properly mapped read pair formation. The black line represents an assembly. The arrows represent all possible read mapping locations. The cases a) and b) correspond to the situations when no read pairs are formed or just one read pair is formed, respectively. The cases c) and d) show examples when several read pairs are formed from two given reads. The case d) is an example of the situation when reads are mapped to a tandem repeat.

Then the created properly mapped read pairs are divided into 4 groups, based on the presence of alternative alignments for each read in a pair:

1. Single group - consisting of pairs where both reads are mapped uniquely to a sequence. The pairs from this group point to the non-repeated regions of a genome (Figure 3a).
2. Single_Multiple group - consisting of pairs where the forward-oriented read is mapped uniquely to a sequence and the reverse-oriented read has multiple alternative mapping locations. The pairs point to the regions where non-repeated regions end and repeated regions start (Figure 3b).
3. Multiple_Single group - consisting of pairs where the forward-oriented read has multiple alternative mapping locations and the reverse-oriented read is mapped uniquely to a sequence. The pairs point to the regions where repeated regions end and non-repeated regions start (Figure 3c).
4. Multiple group - consisting of pairs where both reads have multiple mapping locations. The pairs point to the repeated regions of a genome (Figure 3d).

### 2.3 Read path creation and path gap detection

During the third step, reads from each group are merged together to form continuous paths. This is done separately for forward-and reverse-oriented reads. Only neighbouring reads having an overlap of more than 5 bases are involved in the merging process. There may be small substitution, deletion and insertion differences in reads inside the overlapping regions. If neighbouring reads overlap with 5 or less bases, the overlapped bases are clipped, creating an uncovered region between them. The 5-base limit has been introduced to exclude overlaps appearing due to uncertainties in alignment rather than actual overlaps of positions. The obtained paths represent the fragments of a genome that are considered free of assembly errors.

**Figure 3:**
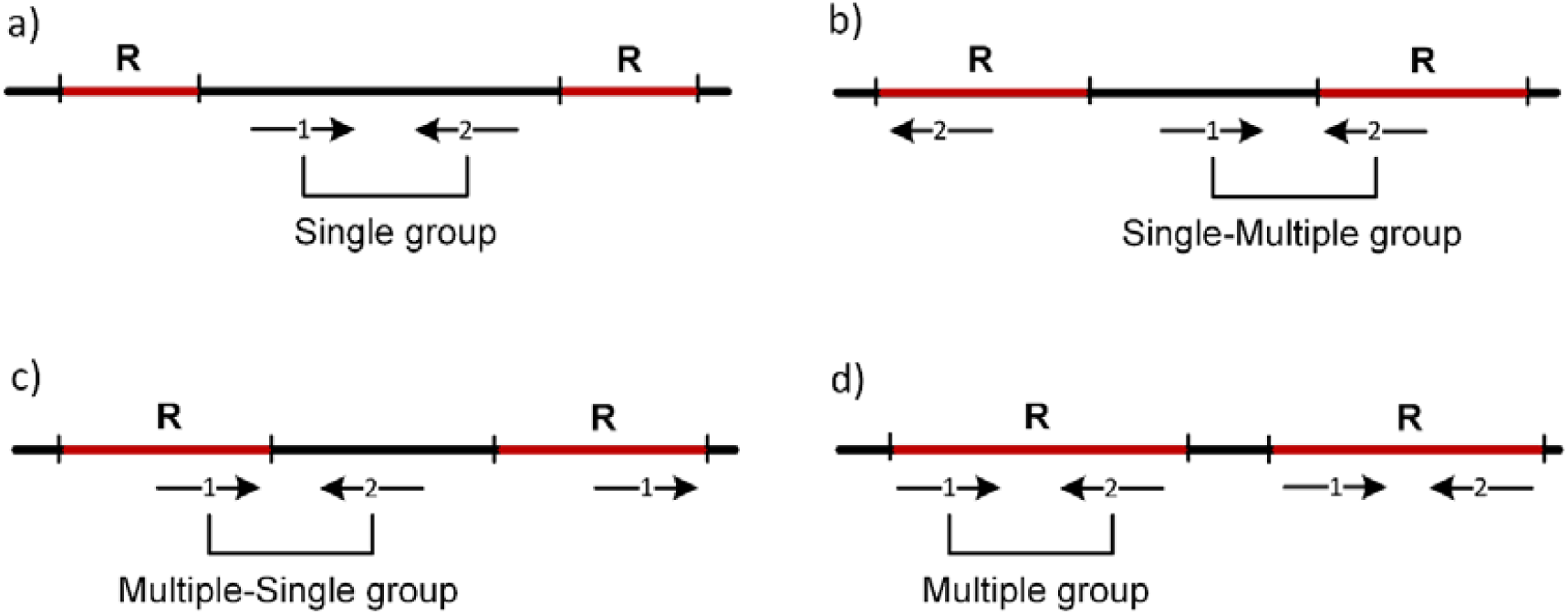
Properly mapped read pair categorization. The black line represents an assembly. The assembly regions marked by red colour correspond to repeated regions. The repeated regions are identical or near-identical copies of the same repeat. The arrows represent all possible read mapping locations.

Usually, several paths of the same type and orientation cover a full assembly sequence. The assembly sequence regions located between paths of the same type and orientation are called path gaps (see Figure 4). The path gaps may potentially contain assembly errors and, therefore, are extensively analysed by NucBreak during the next step.

**Figure 4:**
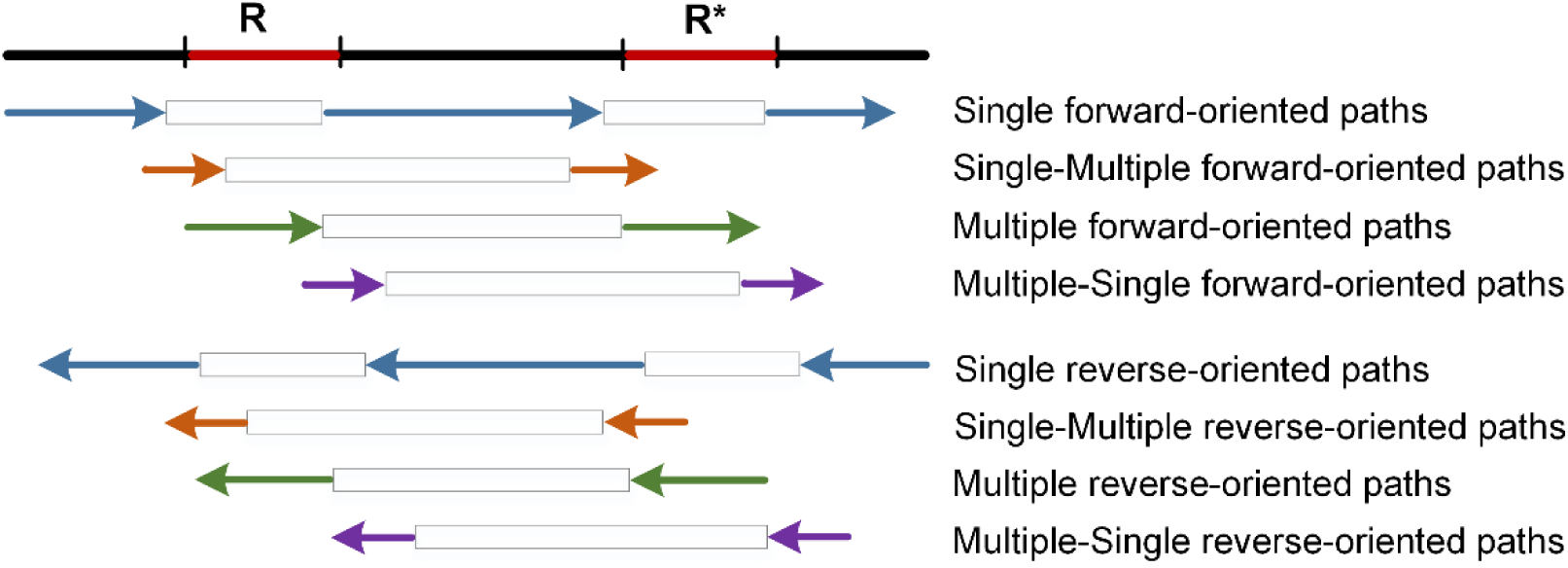
Read paths and path gaps. The black line represents an assembly. The assembly regions marked by red colour correspond to repeated regions. The repeated regions are identical or near-identical copies of the same repeat or copies of different repeats. The arrows represent read paths. The arrows of the same colour correspond to the read paths of the same type. The rectangles between the read paths indicate path gaps. The example demonstrates the correct order of the read paths in the absence of assembly errors.

### 2.4 Path gap analysis

There can be several reasons for path gaps to appear. First, a path gap may appear due to the presence of an assembly error in this region. Second, a path gap may correspond to a region covered by paths of other types. Third, a path gap may appear when there is not enough read coverage to provide the required overlap between reads. Such a situation may occur when: (1) a genome or its fragments were sequenced with a very low coverage, (2) read pairs from these regions are absent due to sequencing errors in reads, (3) read pairs are filtered out due to a violation of condition 4: when there are sequencing errors in the beginning of one of the read in a pair, and (4) there are gaps (a subsequence of N’s) in the genome.

The goal of the fourth step is to exclude path gaps that do not contain assembly errors. NucBreak starts with excluding path gaps that do not overlap with path gaps between paths of the same type in the opposite orientation (Figure 5). Such situations are often observed in Single and Multiple paths and are due to low coverage by either forward-or reverse-oriented reads.

**Figure 5:**
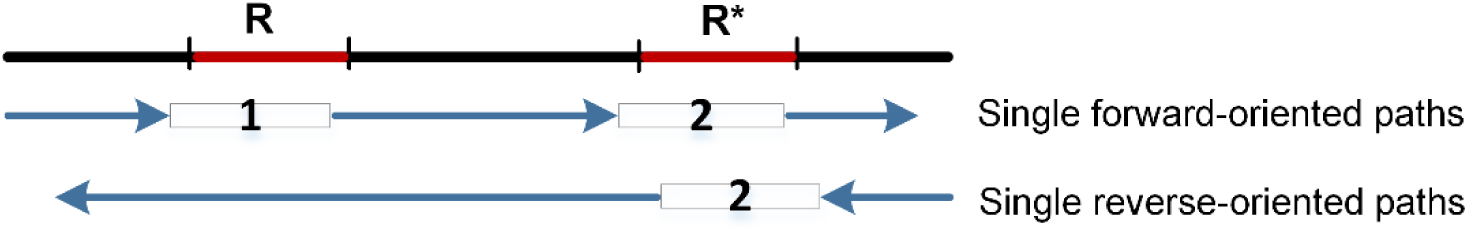
Path gap exclusion. The black line represents an assembly. The assembly regions marked by red colour correspond to repeated regions. The repeated regions are identical or near-identical copies of the same repeat or copies of different repeats. The arrows represent read paths. The rectangles between read paths indicate path gaps. Path gap 1 is excluded because it is fully covered by read path of the same type and another orientation. The path gaps marked by number 2 are not excluded and require further analysis.

Then NucBreak detects path gaps appearing because of the alternation of paths of different types. To accomplish this, NucBreak analyses the location order of path types and the locations of paths separately for the forward-and reverse-oriented paths. The path types should appear in a fixed order, like a cycle: Single, Single_Multiple, Multiple, Multiple_Single, Single, and so on (Figure 4). The cycle may start with any type. If one type is skipped or repeated (Figure 6), it indicates an error in this region. There is also a requirement for the locations of paths: both a path and the following path gap should overlap with the next path with more than 5 bases. However, we make some exceptions for type order and path locations in special cases (see [Additional file 1: Figure S1] for the details). In this way, NucBreak excludes a path gap if the beginning of the path gap is covered with a path that has a correct type order and location.

**Figure 6:**
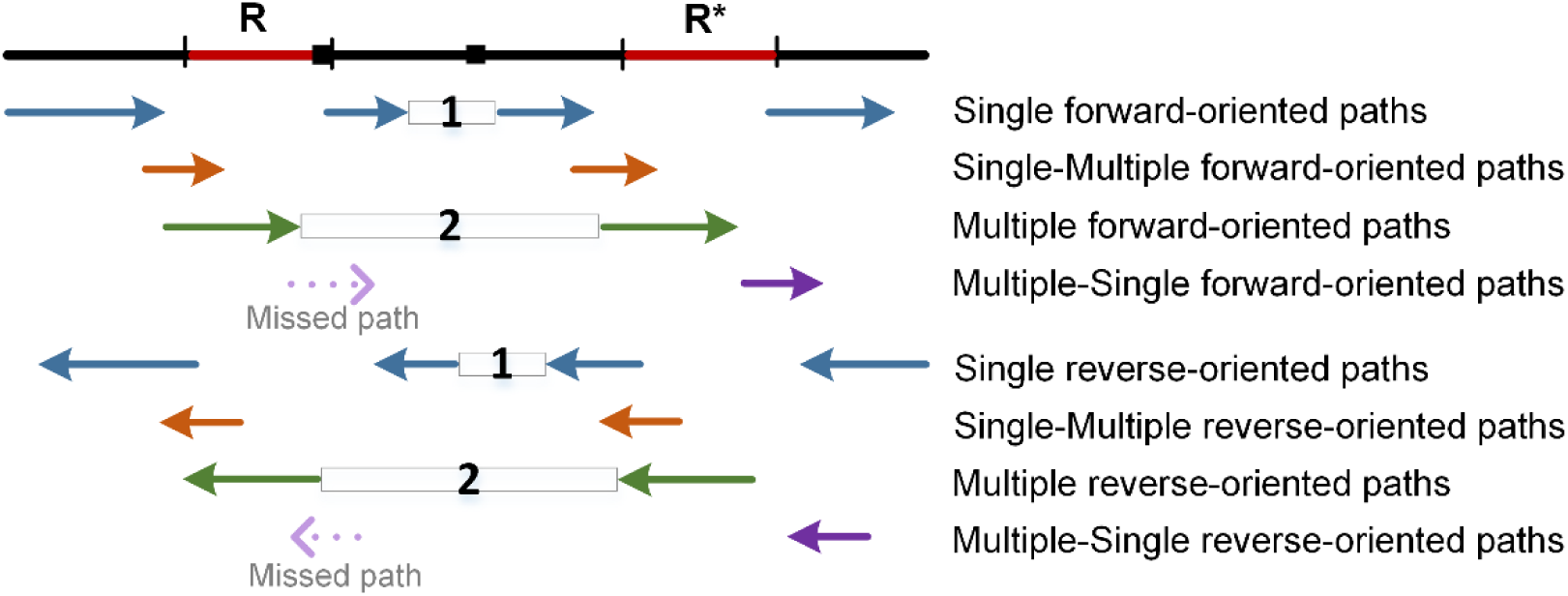
Path gap exclusion. The black line represents an assembly. The assembly regions marked by red colour correspond to repeated regions. The repeated regions are identical or near-identical copies of the same repeat or copies of different repeats. The black squares mark the locations of assembly errors. The arrows represent read paths. The rectangles between read paths indicate path gaps that are not excluded. The path gaps marked by number 1 is not excluded due to the repetition of read path types (e.g. the Single forward-oriented path is followed by another Single forward-oriented path instead of the Single-Multiple forward-oriented path). The path gaps marked by number 2 are not excluded because one read path type is missed (e.g. Multiple forward-oriented path is followed by Single forward-oriented path instead of Multiple-Single forward-oriented path).

Unfortunately, it is not always possible to exclude all path gaps located in the assembly sequence regions that do not contain errors. The path gaps that have appeared due to low read coverage or are located in the regions containing subsequences of N’s of appropriate lengths are never excluded.

### 2.5 Error location adjustment

All non-excluded path gaps are treated as containing assembly errors. To narrow down the region where an error is located, NucBreak shortens the path gaps during the fifth step. To accomplish this, it first combines the paths of all types with the same direction together. Then for each path gap, it determines whether the end of any path is inside the path gap region. If it is, the path gap beginning is shifted to the path end (or to the right-most end in case of several paths detected, Figure 7a). Finally, it determines whether the beginning of any path is inside the path gap region. If it is, the path gap end is shifted to the path beginning (or to the left-most beginning in case of several paths detected, Figure 7b). If any path gap is fully covered by any path, then this path gap is excluded.

**Figure 7:**
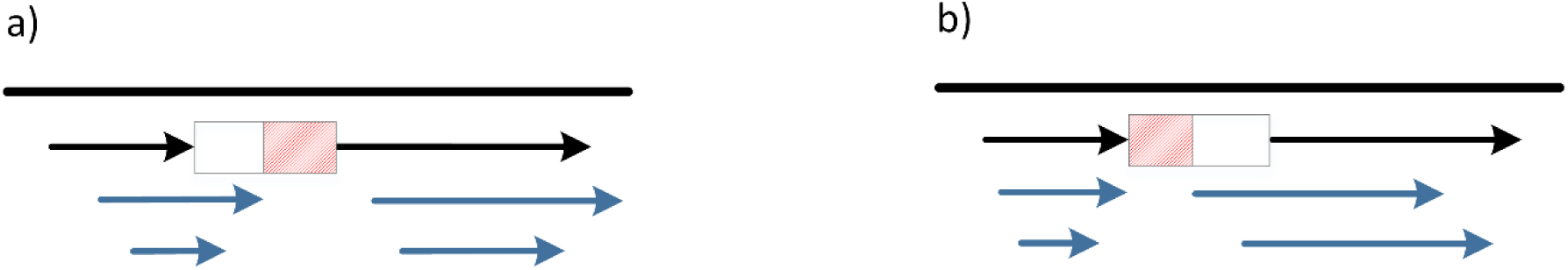
Error location adjustment. The black line represents an assembly. The arrows represent read paths of any type. The rectangles represent initial path gaps. The red areas in the rectangles in cases a) and b) correspond to the adjusted path gaps with the shortened beginning and end, respectively.

To pinpoint the locations of errors, NucBreak first finds the union of the adjusted path gaps of all types. This is carried out separately for path gaps located on forward-and reverse-oriented paths. Then NucBreak finds the intersection of the obtained forward-and reverse-oriented unions of regions and pinpoints the error locations. Errors in the beginning and at the end of a sequence (inside the regions with lengths equal to the read length) are excluded by NucBreak, because in most cases they are due to the lack of perfectly mapped read pairs.

### 2.6 Data sets

For the testing purposes, we created four different datasets. For the first and second datasets, we constructed artificial reference genomes and assemblies, and generated simulated Illumina paired-end read libraries. In both datasets, the reference genomes were constructed from random DNA sequences by introducing different interspersed and tandem repeats. The assemblies were generated from the reference genomes sequences by introducing controlled modifications (e.g. relocations, deletions, duplications of different fragments and so on). The detailed description of introduced modifications is given in [Additional file 1: Table S1]. Depending on the datasets, different approaches were applied to create an Illumina paired-end read library in each case. For the first dataset, one read library was generated with the help of ART (Q version 2.5.8) [9] run with the “-ss MSv3 -l 250 -p -m 700 -s 40” settings with 40x read coverage for each reference genome. For the second dataset, read libraries with 5x,10x, 40x,100x, and 200x read coverages were generated by ART run with the “-ss MSv3 -l 250 -p -m 700 -s 40” settings.

The third dataset was created on the base of the data provided by the Assemblathon 1 project [10]. An artificially evolved human chromosome 13 (hg18/NCBI36), simulated Illumina paired-end read library with 40x coverage, and genome assembly obtained by PE-assembler [11] were downloaded from the Assemblathon 1 website [12]. To increase the number of errors and to introduce more variability of error types, we deleted all gaps from the assembly.

The fourth dataset consisted of 8 bacterial genomes (*Bordetella pertussis* str. J081, *Brucella melitensis* str. 1, *Enterobacter cloacae* str. AR_0136, *Escherichia coli* str. 2014C-3599, *Klebsiella pneumoniae* str. SGH10, *Pseudomonas aeruginosa* str. AR_0095, *Salmonella enterica* str. CFSAN047866, and *Staphylococcus aureus* str. CFSAN007896), MiSeq Illumina paired-end reads libraries provided for these genomes, and assemblies generated using the ABySS (version 2.0.2) [13], SPAdes (version 3.11.0) [14] and Velvet (version 1.2.10) [15] assemblers. The genomes were downloaded from the NCBI database [16], and the reads were downloaded from the EBI database [17]. The genomes accession numbers and information about the read libraries are given in [Additional file 1: Table S2]. The parameter settings used to run ABySS, SPAdes and Velvet are described in [Additional file 1]. As in the third dataset, we have removed all gaps from the assemblies before testing.

## 3. Results

We have created a tool called NucBreak that is aimed at detection of structural errors in assemblies by analysing the placements of properly mapped reads and exploiting information about the alternative alignments of the reads. In this section, we examine the ability of NucBreak as well as REAPR (version 1.0.18), FRCbam (version 1.2.0), Pilon (version 1.22), BreakDancer (version 1.3.6), Lumpy (version 0.2.13), and Wham (version 1.8.0) to detect assembly errors in real and simulated datasets.

All tools, except REAPR, FRCbam and partly NucBreak, were run with their default settings. The parameter settings used to run REAPR, FRCbam and NucBreak are described in [Additional file 1]. To validate the results, we compared the obtained results of each test with the ground truth results consisting of real errors. Depending on the test performed, the ground truth results were generated during the simulation process or produced using NucDiff [18], the tool which enables comparison of reference genomes with assemblies. NucDiff was run with the default parameter settings. The ground truth and obtained results were compared using BEDTools (version 2.17.0) [19] to get sensitivity and precision for each tool and each dataset (see [Additional file 1] for more details).

We studied sensitivity and precision for each tool allowing various degrees of slack in the location of each region in the comparison by adding flanking regions of different sizes to the ground truth regions. We added 1, 5, 10, 20, 50, 100, 200, 400, and 600 bp both up-and downstream of each ground truth entry. The flanking regions were introduced to investigate the positional accuracy of the tools tested. In addition, for the flanking region size equal to 600 bp, we identified the ability of each tool to detect ground truth errors depending on the read coverage value in one of the tests.

### 3.1 Accuracy assessment in simulated datasets

We created a simulated dataset consisting of ten artificial reference genomes, assemblies, and Illumina paired-end read libraries, as described in Section 2.6 (the first dataset), and ran NucBreak, Pilon, REAPR, FRCbam Lumpy, Wham and BreakDancer to detect errors in the assemblies. To enable validation of the obtained results, we also generated the ground truth results during the simulation process. All ground truth errors were divided into several groups according to their types and sizes: insertion, duplication, tandem duplication, deletion, deletion of interspersed repeats or their parts, deletion of tandem repeats or their parts, inversion, relocation (intra-chromosomal rearrangements) with either inserted regions between misjoined regions or without them, and relocation with overlapped misjoined regions groups with error sizes between 10 and 49 bp, 50 and 299 bp, and greater than 299 bp. We also categorized the obtained errors into four basic groups: deletion, insertion, inversion and relocation groups. The relations of obtained errors with the repeated sequence regions as well as their sizes were not taken into account. The obtained error type categorization became possible due to a specific way of assembly modification during the simulation process. The sensitivity and precision results are presented in Figures 8-11. The number of ground truth errors in each group is given in [Additional file 1: Table S3]

**Figure 8:**
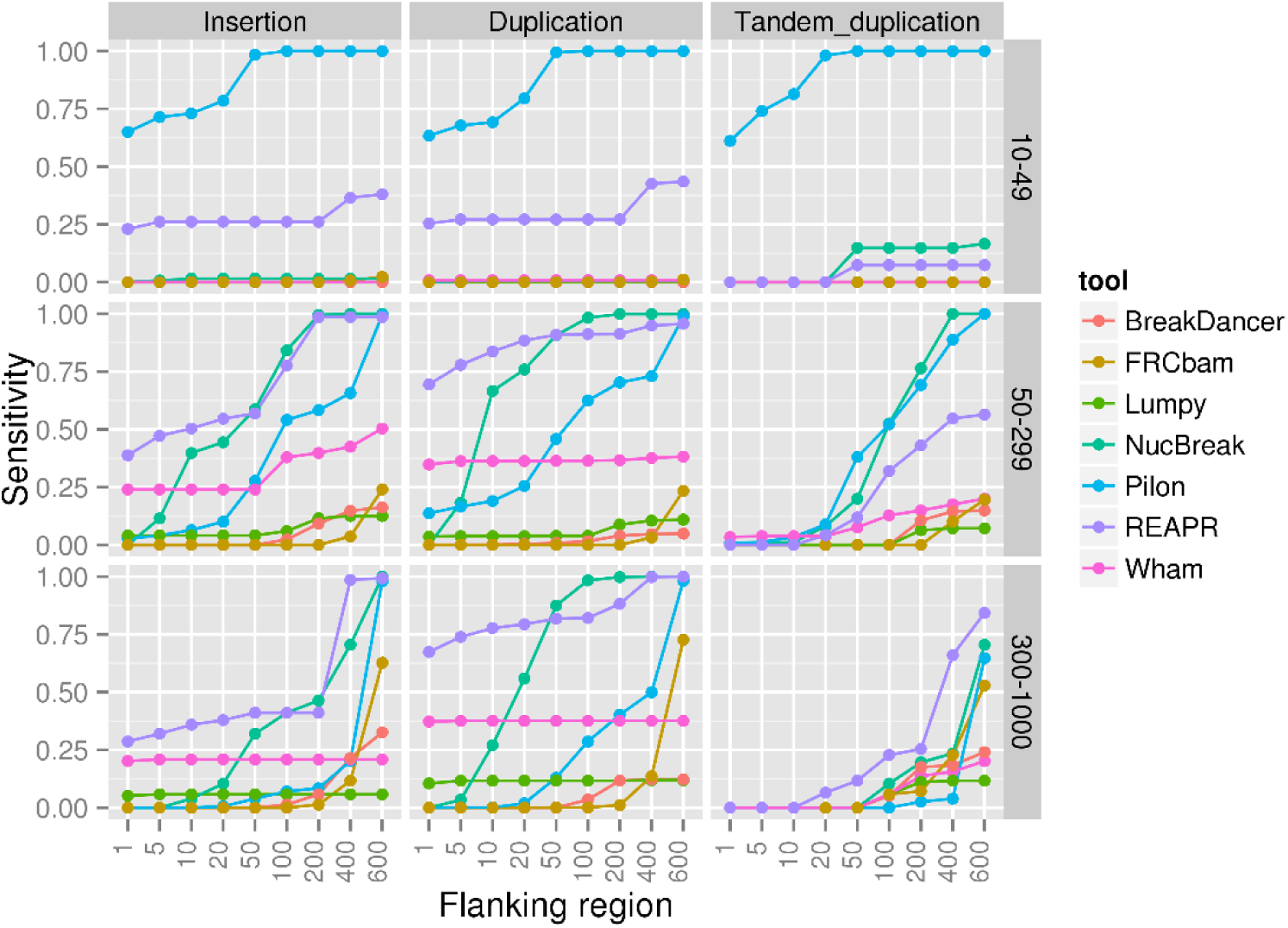
Sensitivity results for the insertion, duplication and tandem duplication groups, obtained using the simulated datasets.

**Figure 9:**
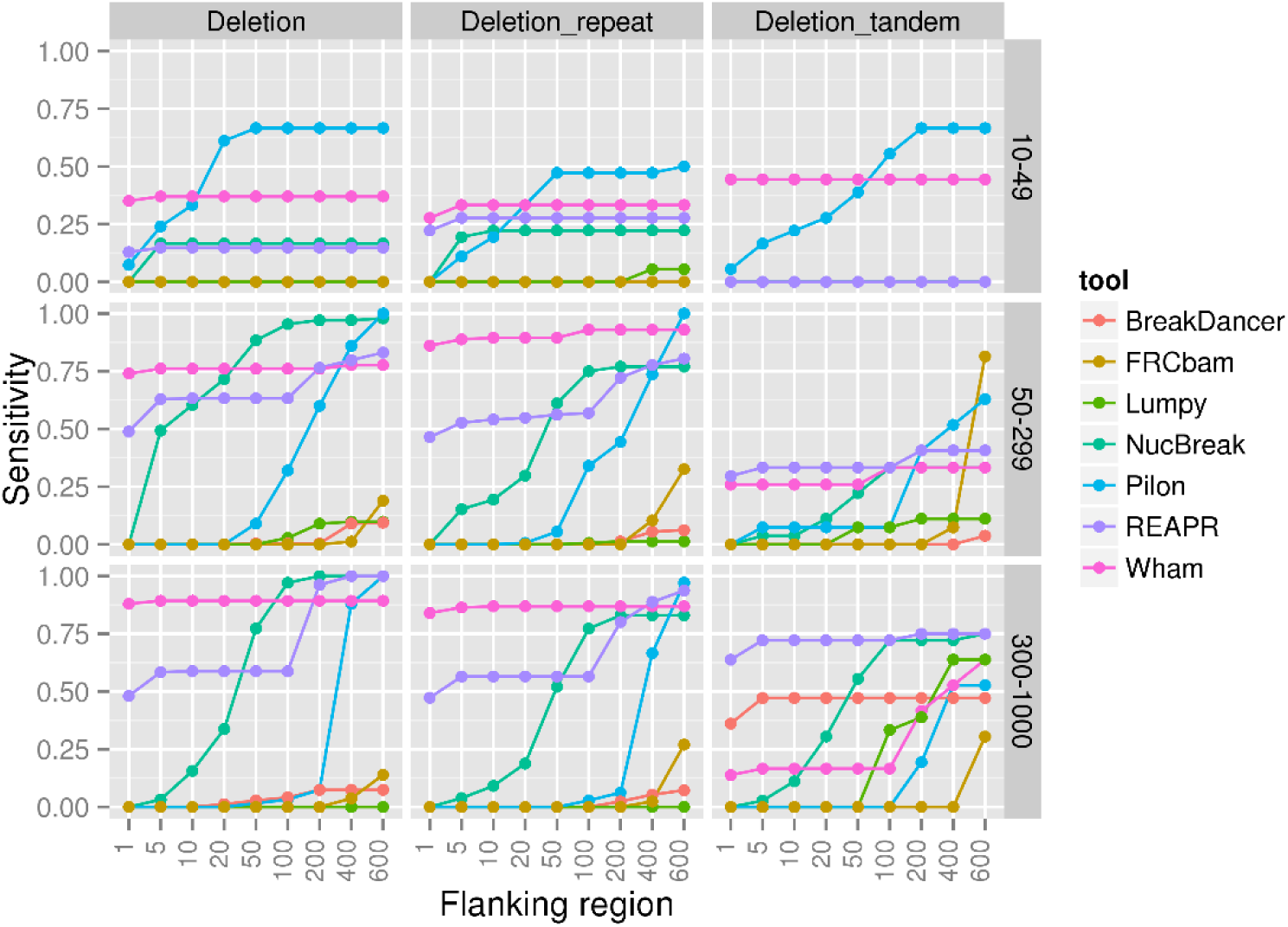
Sensitivity results for the deletion, deletion_repeat and deletion_tandem groups, obtained using the simulated datasets. The deletion_repeat group contains deletions of interspersed repeats or their parts. The deletion_tandem group contains deletions of tandem repeats or their parts.

**Figure 10:**
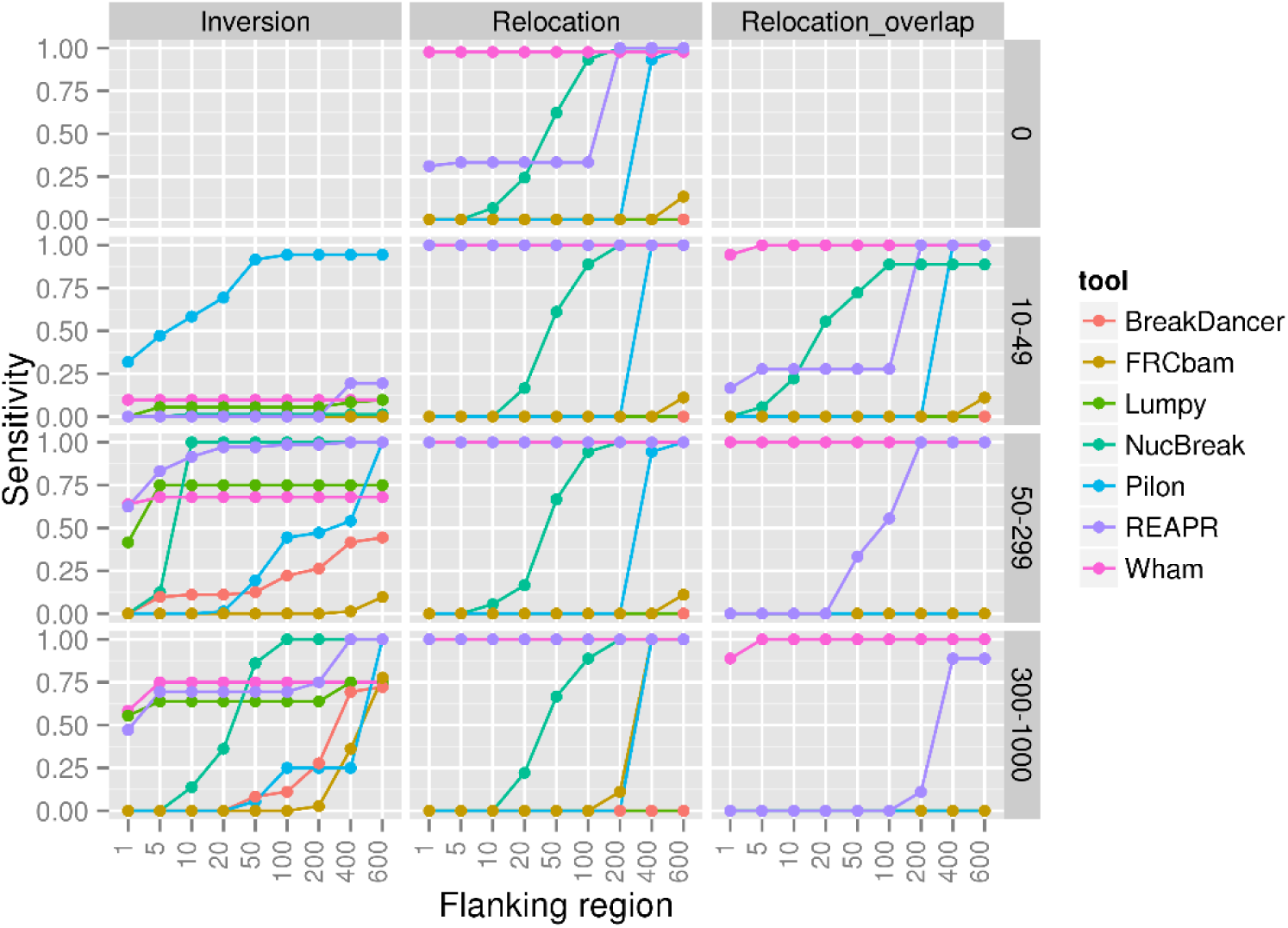
Sensitivity results for the inversion, relocation and relocation_overlap groups, obtained using the simulated datasets. The relocation group consists of relocations with either inserted regions between misjoined regions (size varied between 10 and 1000) or without them (size is equal to 0). The relocation_overlap group consists of relocations with overlapped misjoined regions.

**Figure 11:**
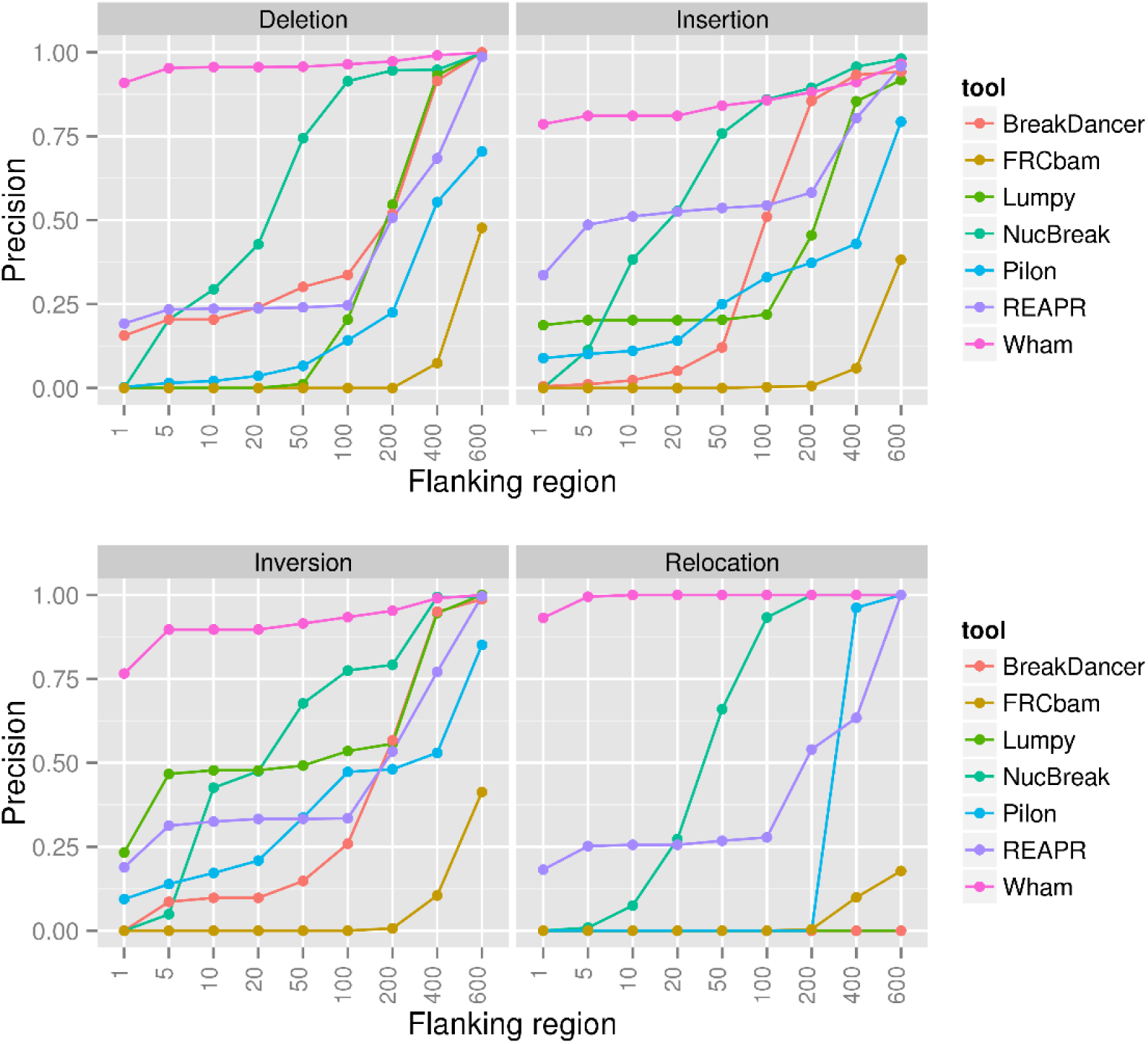
Precision results for the deletion, insertion, inversion and relocation groups, obtained using the simulated datasets.

As can be seen from Figures 8-10, sensitivity of each tool largely depends on the types and sizes of errors and size of the flanking region. For Pilon and NucBreak, the sensitivity constantly increases with respect to flanking region size increment in all cases where sensitivity is larger than zero. Wham’s and REAPR’s sensitivity either increases with respect to the flanking region size increase or remains approximately the same, depending on the error types and sizes. In case of BreakDancer, FRCbam and Lumpy, sensitivity increases starting from medium-or long-sized flanking regions depending on an error group.

As expected, all tools perform best with 600 bp flanking region. For this flanking region size, Pilon obtains sensitivity equal to 1 in almost all error groups and outperforms other tools in many cases. NucBreak’s and REAPR’s sensitivity is the same or close to Pilon’s one in most groups. Wham shows relatively high sensitivity in many groups, while BreakDancer, FRCbam and Lumpy have low sensitivity in almost all cases.

Precision increases slowly for Wham and Lumpy and rapidly for all other tools together with the flanking region size increase. All tools except Pilon and FRCbam reach precision equal to 1 with 600 bp flanking region in all groups except the Relocation group. In the Relocation group, only Wham, NucBreak, Pilon, and REAPR get precision equal to 1.

### 3.2 Accuracy assessment in simulated datasets depending on read coverage

To explore the influence of read coverage on the results of NucBreak, Pilon, FRCbam, REAPR, Wham, Lumpy, and BreakDancer, we created ten simulated reference genomes, assemblies, and Illumina paired-end read libraries with 5x, 10x, 40x, 100x, and 200x coverage as described in the Section 2.6 (the second dataset). As well as in the Section 3.1, the ground truth errors were generated during simulation process and divided into different groups based on the error types and size. The obtained errors were also divided into deletion, insertion, inversion and relocation groups. The sensitivity and precision values were calculated with 600 bp flanking region. The 600 bp flanking region was chosen because all tools performed best with this flanking region size in the previous section. The sensitivity results are presented in Figures 12-14 and the precision results are shown in Figure 15.

**Figure 12:**
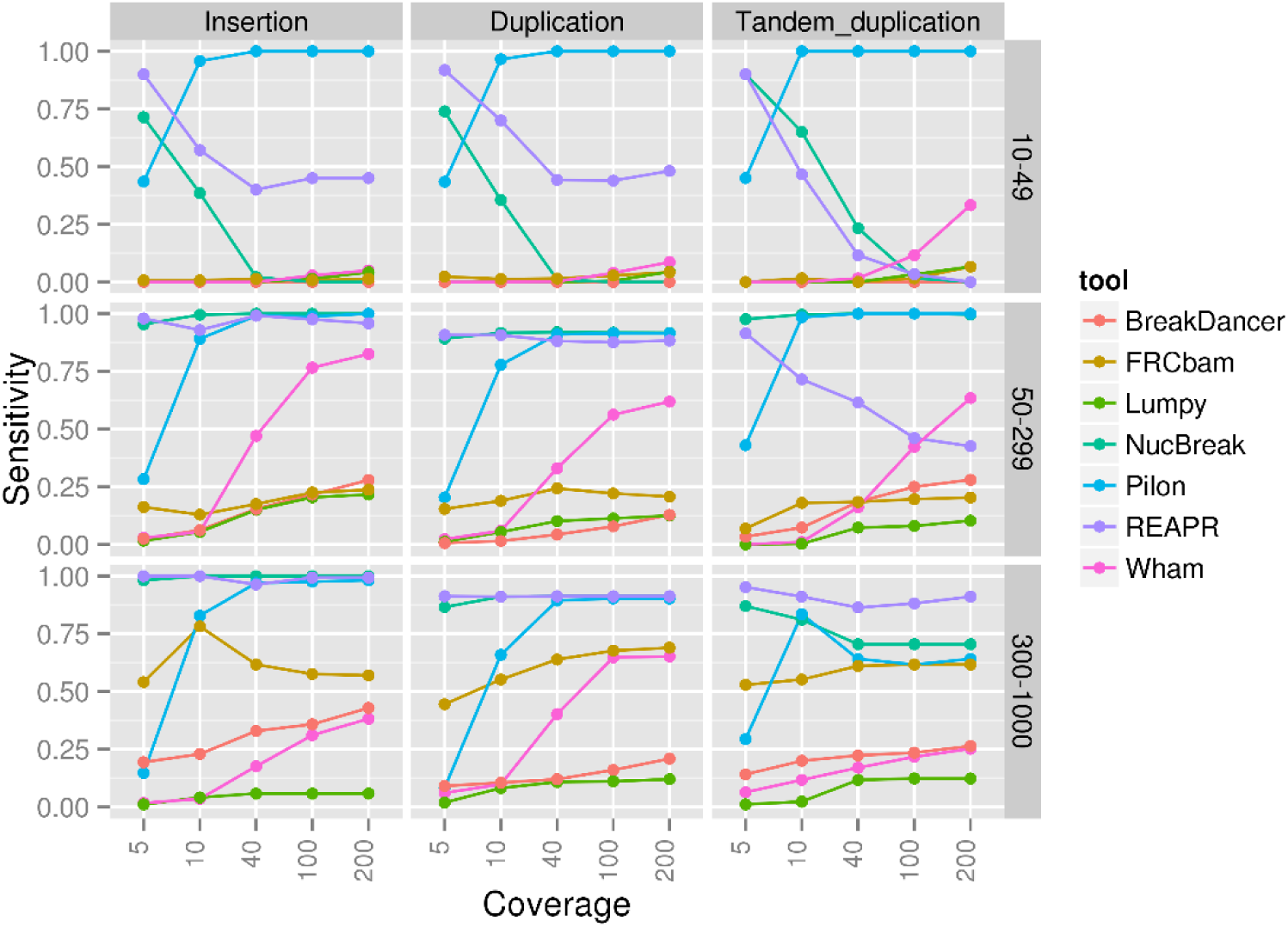
Sensitivity results for the insertion, duplication and tandem duplication groups, obtained using the simulated datasets.

**Figure 13:**
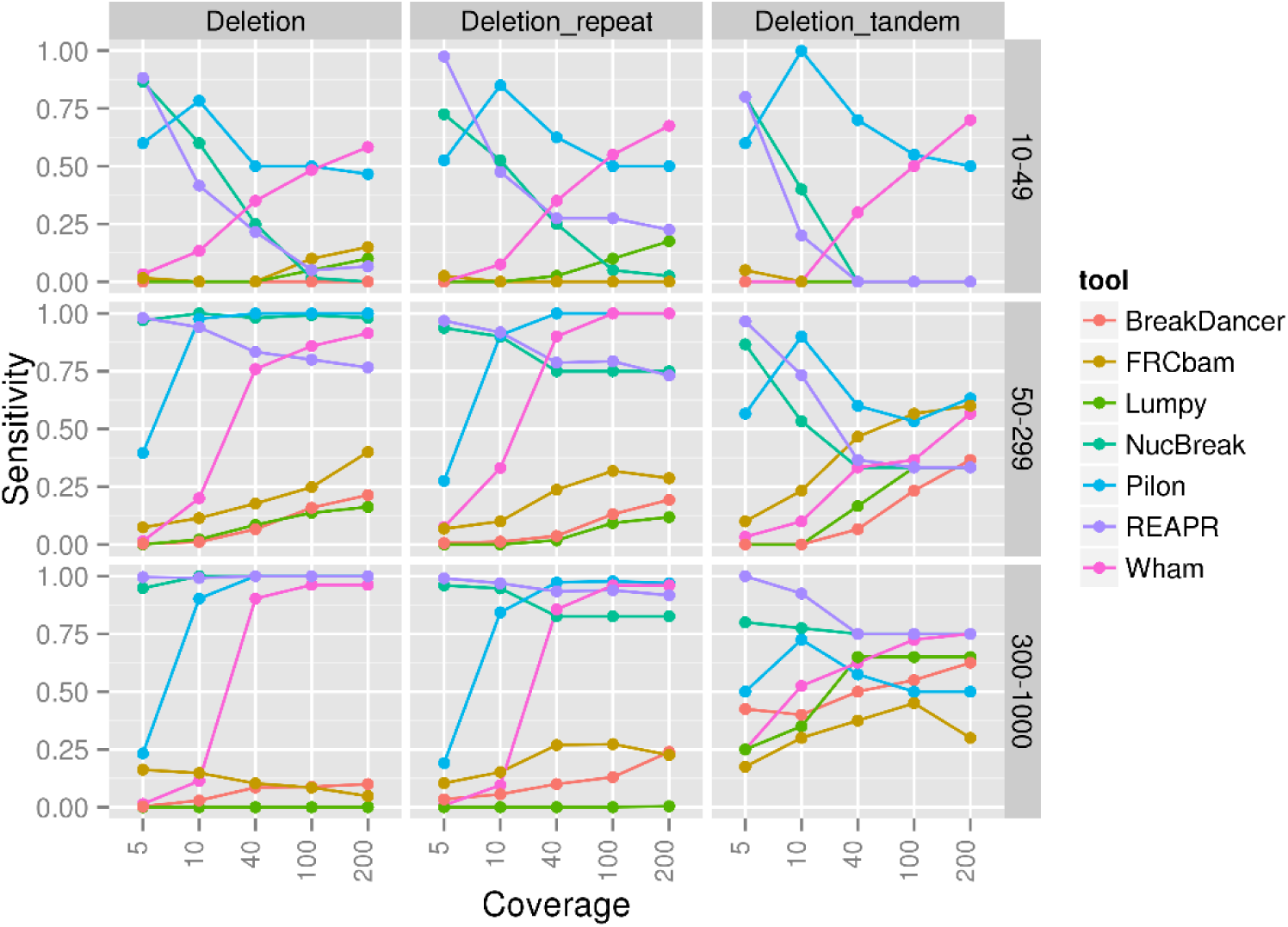
Sensitivity results for the deletion, deletion_repeat and deletion_tandem groups, obtained using the simulated datasets. The deletion_repeat group contains deletions of interspersed repeats or their parts. The deletion_tandem group contains deletions of tandem repeats or their parts.

**Figure 14:**
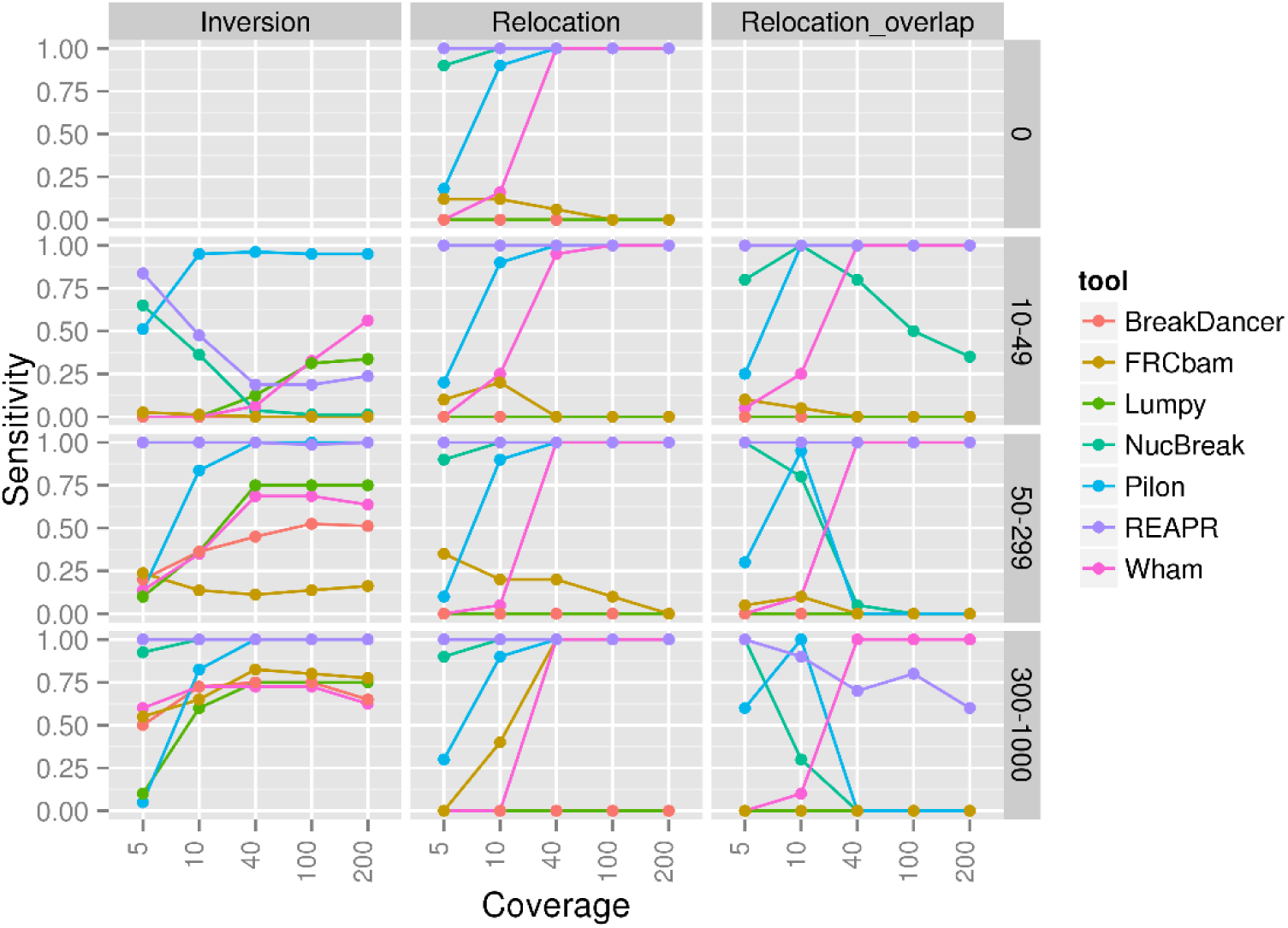
Sensitivity results for the inversion, relocation and relocation_overlap groups, obtained using the simulated datasets. The relocation group consists of relocations with either inserted regions between misjoined regions (size varied between 10 and 1000) or without them (size is equal to 0). The relocation_overlap group consists of relocations with overlapped misjoined regions.

**Figure 15:**
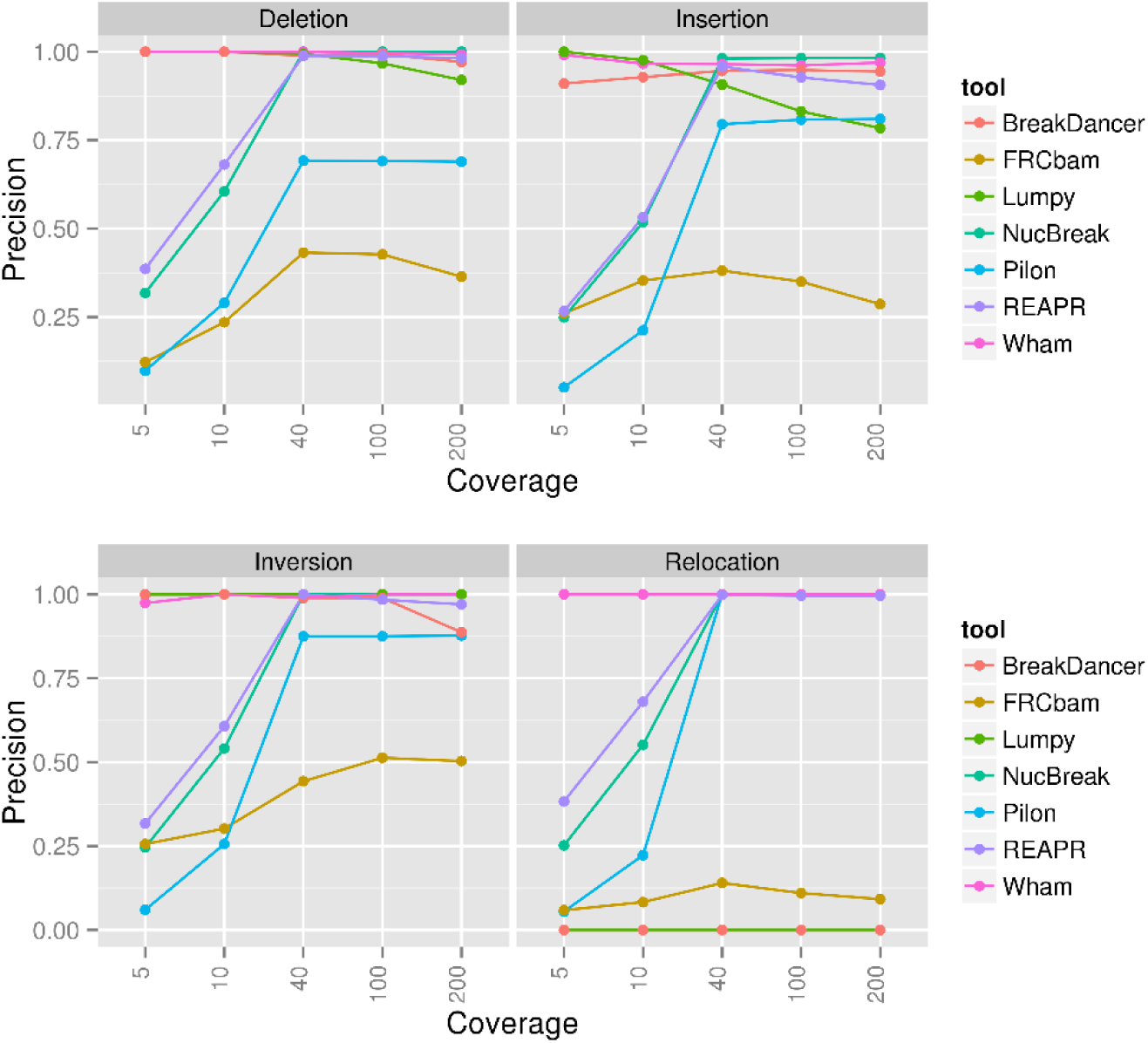
Precision results for the deletion, insertion, inversion and relocation groups, obtained using the simulated datasets.

As indicated in provided plots, NucBreak’s and REAPR’s sensitivity either decreases with the coverage increase or is approximately the same starting from 10x coverage. In case of NucBreak, the sensitivity decrease can be explained by the fact that the number of regions where reads are not overlapped are reduced with increased coverage, and, thus, less assembly errors are predicted just by chance. Pilon’s and FRCbam’s sensitivity decreases or increases depending on the error type and coverage values, while in case of Wham, BreakDancer, and Lumpy sensitivity always increases, except a small number of cases when the sensitivity remains approximately the same.

REAPR, NucBreak and Pilon demonstrate the fast increase of precision in all groups with up to 40x coverage. Starting from 40x coverage, precision remains the same or slightly decreases. In case of Lumpy, FRCbam, BreakDancer and Wham, precision remains approximately the same for all coverage values or slightly changes with coverage increasing.

### 3.3 Accuracy assessment in an assembly obtained from simulated reads

To validate the ability of NucBreak, Pilon, REAPR, FRCbam, Lumpy, BreakDancer, and Wham to detect errors in real assemblies, we ran the tools with a dataset where reads were created for an artificially evolved diploid genome and an assembly was generated by the PE-assembler (see Section 2.6, the third dataset for details). The ground truth results were obtained by comparing the assembly with the reference genome using NucDiff. All ground truth errors were divided into types according to the error types and sizes provided by NucDiff: substitution, insertion, duplication, tandem duplication, deletion, deletion of interspersed repeats or their parts, deletion of tandem repeats or their parts, inversion, reshuffling (several neighbouring genome regions are placed in a different order in an assembly), and two groups of rearrangements (arrangement and rearrangement with overlap) with sizes between 10 and 49 bp, between 50 and 299 bp, and greater than 299 bp. The rearrangement group consisted of relocation and translocation (an inter-chromosomal rearrangement) errors with either inserted regions between misjoined regions or without them. The relocation with overlap group contained relocation and translocation errors with overlapped misjoined regions. The obtained results were not grouped because not all tested tools reported types of detected errors, and, thus, total precision was calculated for each tool. The sensitivity and precision results are presented in Figures 16-20. The number of ground truth errors in each group is given in [Additional file1: TableS3].

**Figure 16:**
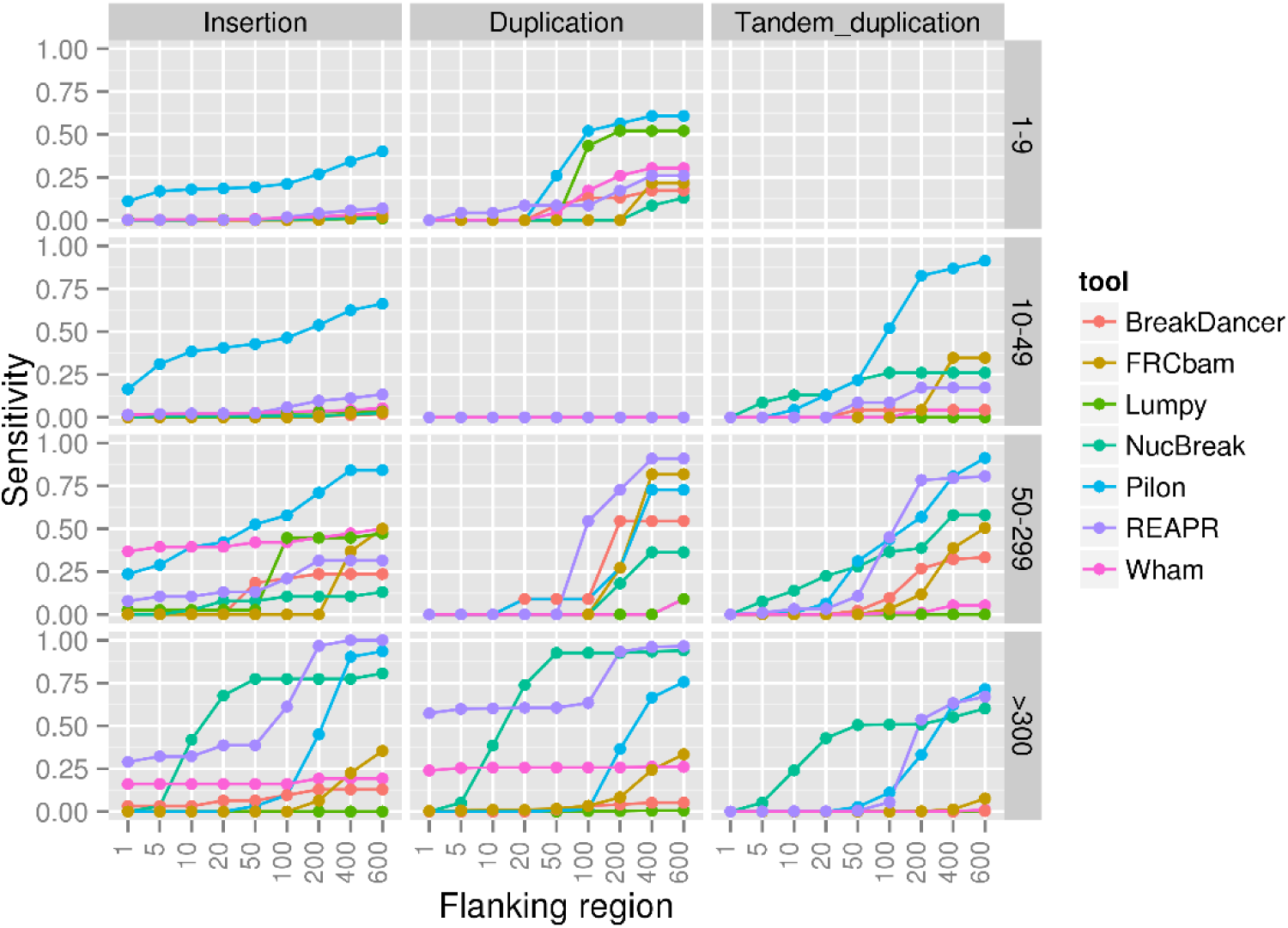
Sensitivity results for the insertion, duplication and tandem duplication groups, obtained using the datasets from the Assemblathon 1 project.

**Figure 17:**
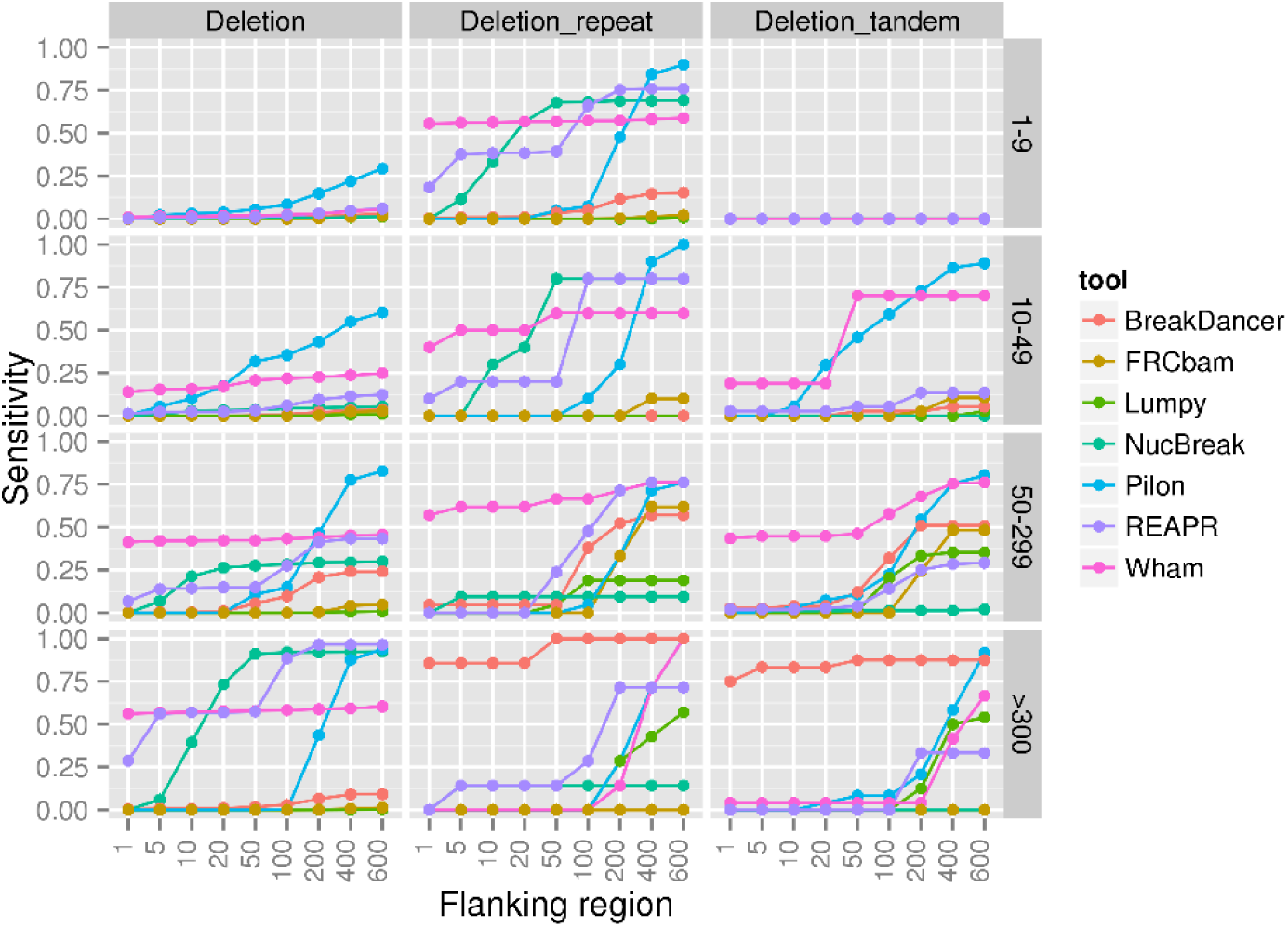
Sensitivity results for the deletion, deletion_repeat and deletion_tandem groups, obtained using the datasets from the Assemblathon 1 project. The deletion_repeat group contains deletions of interspersed repeats or their parts. The deletion_tandem group contains deletions of tandem repeats or their parts.

**Figure 18:**
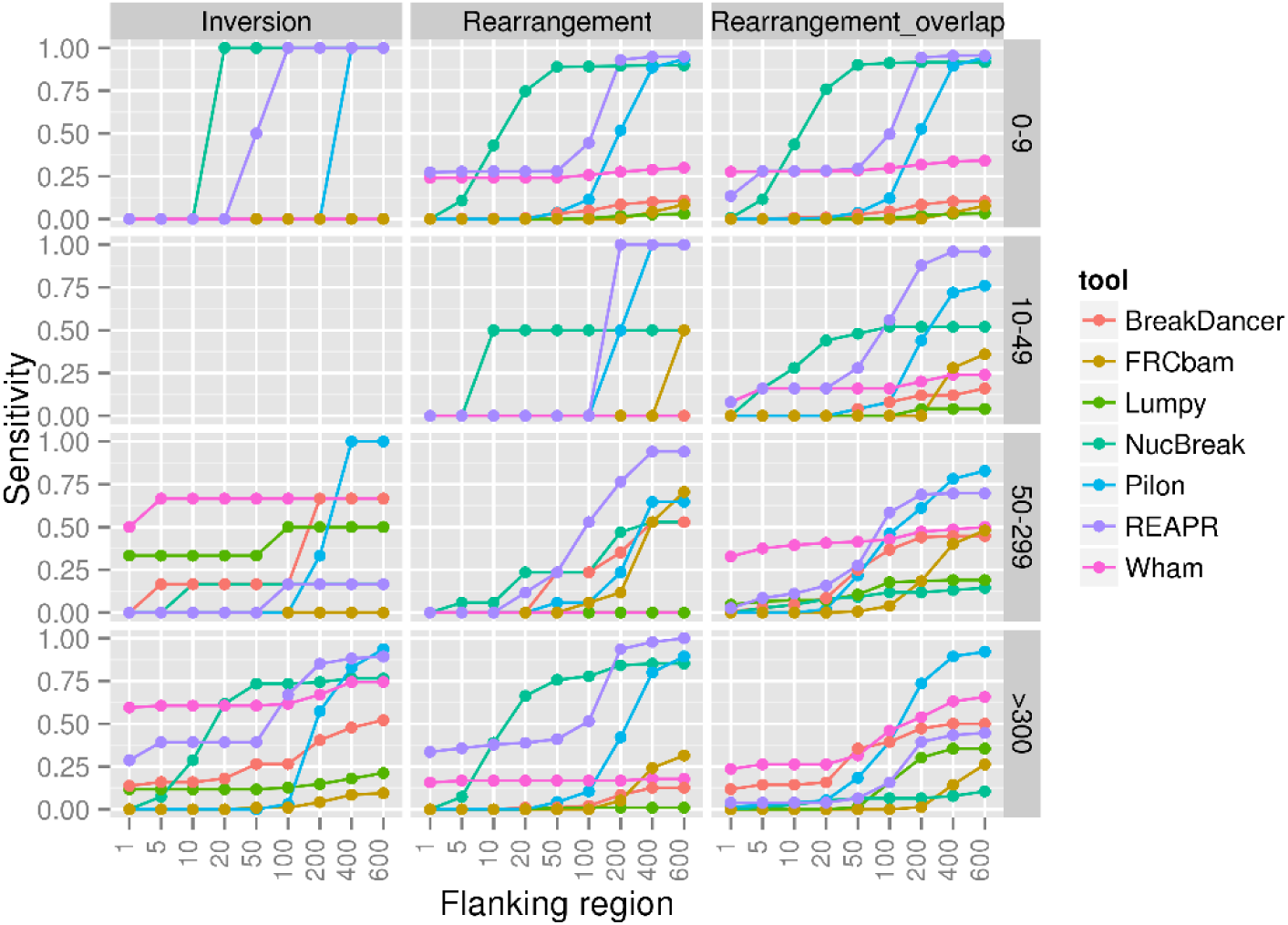
Sensitivity results for the inversion, rearrangement and rearrangement_overlap groups, obtained using the datasets from the Assemblathon 1 project. The rearrangement group consists of relocations and translocations with either inserted regions between misjoined regions (size varied between 1 and 1000) or without them (size is equal to 0). The rearrangement_overlap group consists of relocations and translocations with overlapped misjoined regions.

**Figure 19:**
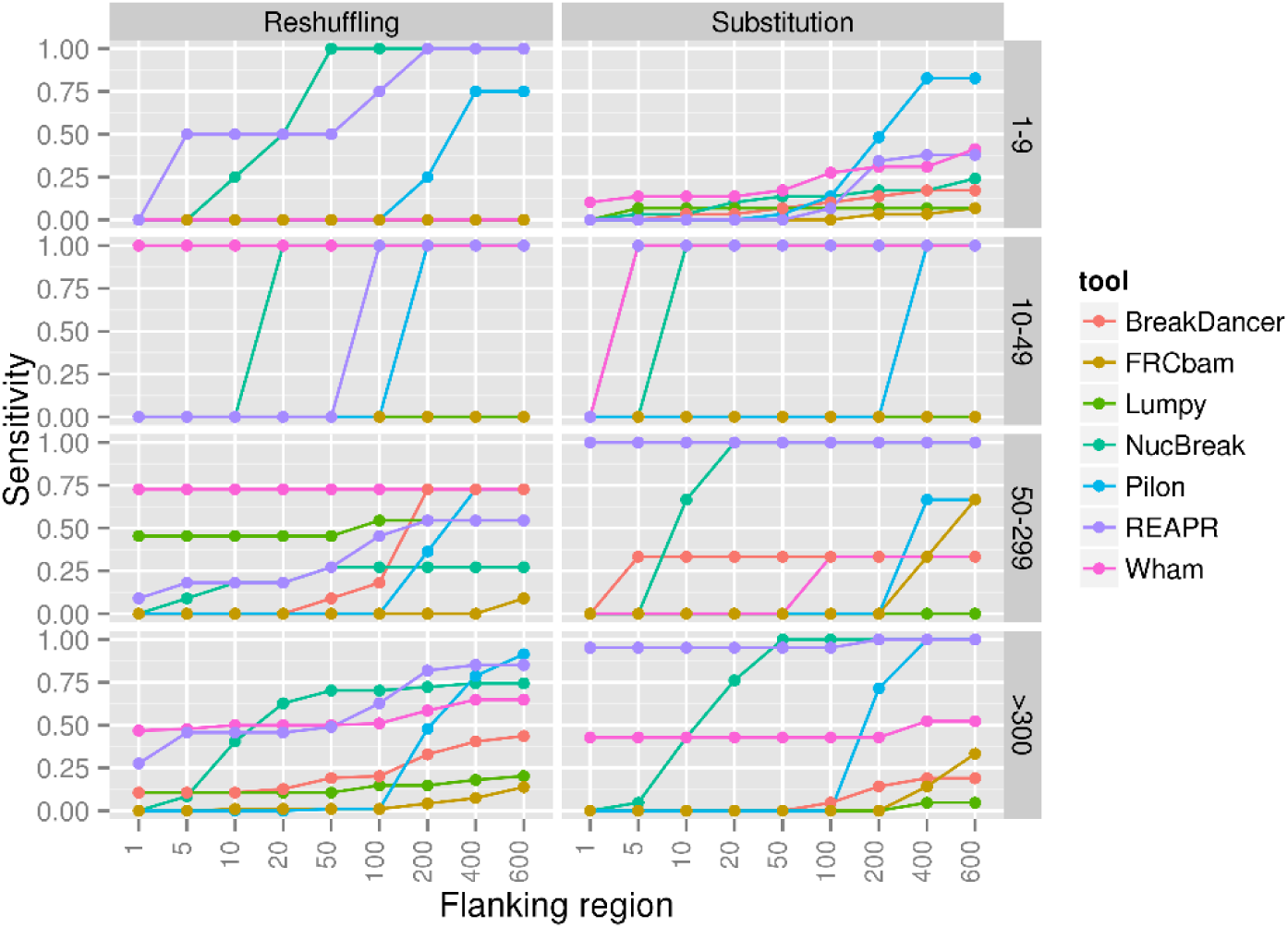
Sensitivity results for the reshuffling and substitution groups, obtained using the datasets from the Assemblathon 1 project.

**Figure 20:**
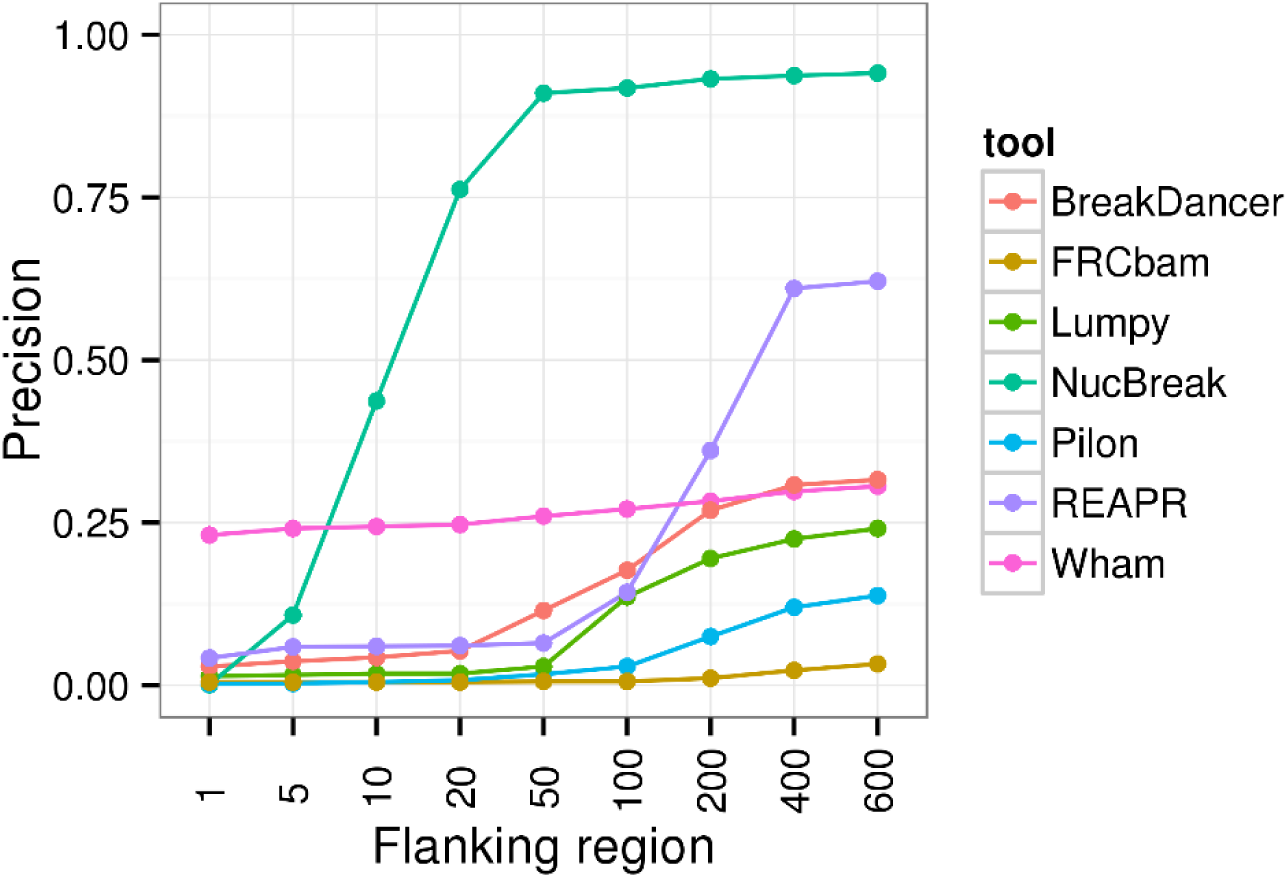
Overall precision obtained using the datasets from the Assemblathon 1 project.

As we see from Figures 16-20, the sensitivity and precision increase with flanking region size increment for all tools in all groups. As expected, all tools perform best with 600 bp flanking region. For this flanking region size, Pilon shows high sensitivity in almost all error groups and outperforms other tools in many cases. The sensitivity results of the other tools largely depend on types and sizes of detected errors. However, all tools show high sensitivity in some groups. The precision between tools varies widely. The best precision is obtained by NucBreak and is equal to 0.94. REAPR has the second-best precision, which is 0.62. Precision of all other tools is less than 0.32. When using smaller flanking regions, e.g. 50 bp, NucBreak’s precision is clearly superior to the other tools.

### 3.4 Accuracy assessment in an assembly obtained from real reads

We also explored the ability of NucBreak, Pilon, REAPR, FRCbam, Lumpy, BreakDancer to detect errors in assemblies obtained from real reads. For this purpose, we downloaded reads for eight bacterial genomes, generated assemblies by using Abyss, SPAdes, and Velvet (see Section 2.6 for full description of data and assembler parameter settings used) and ran NucBreak, Pilon, REAPR, FRCbam, Lumpy, BreakDancer, and Wham. Unfortunately, REAPR crashed during execution and was therefore eliminated from the evaluation process. The ground truth errors were obtained by comparison of assemblies with the reference genomes by using NucDiff and categorized into several types according to the error types and sizes provided by NucDiff, in the same way as it was described in Section 3.3. The sensitivity and precision results were first computed separately for each assembly and genome and then combined together, resulting in the overall sensitivity and precision results. The final results are presented in Figures 21-25. The number of ground truth errors in each group is given in [Additional file1: Table S3].

The sensitivity results indicate that Pilon and NucBreak (with some small exceptions) enable detection of ground truth errors in all non-empty groups, and other tools predict errors only in some cases. Pilon outperforms other tools in almost all groups with respect to sensitivity. However, in half of the cases, the NucBreak results are comparable to Pilon’s ones. Pilon and NucBreak have relatively high sensitivity in many cases, while sensitivity of other tools, except Wham’s and FRCbam’s sensitivity in one case, is quite low or equal to 0. The precision is very low for all tools except NucBreak. NucBreak has relatively high precision, even with short flanking regions.

**Figure 21.**
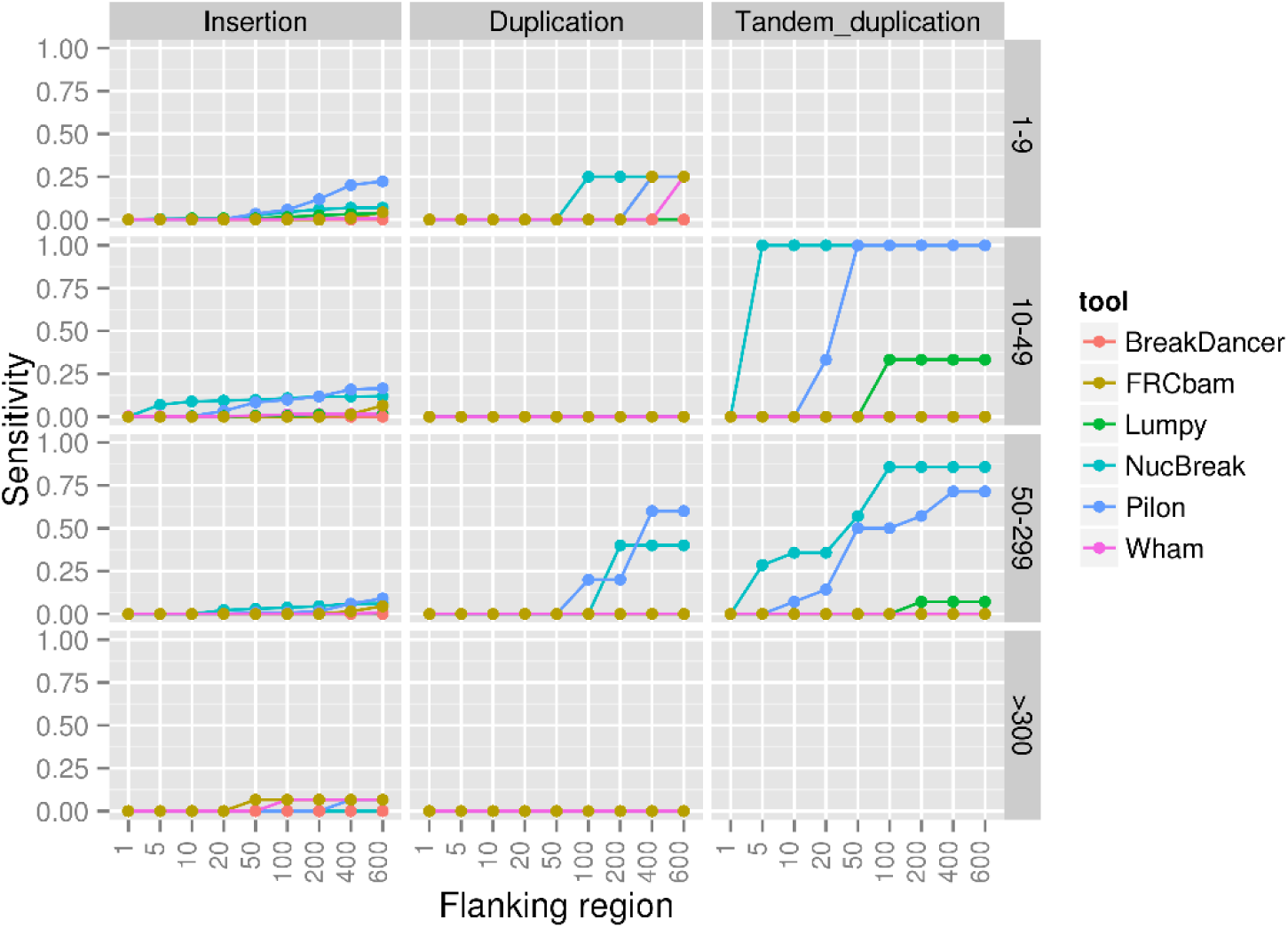
Sensitivity results for the insertion, duplication and tandem duplication groups obtained using the bacterial genome datasets.

**Figure 22.**
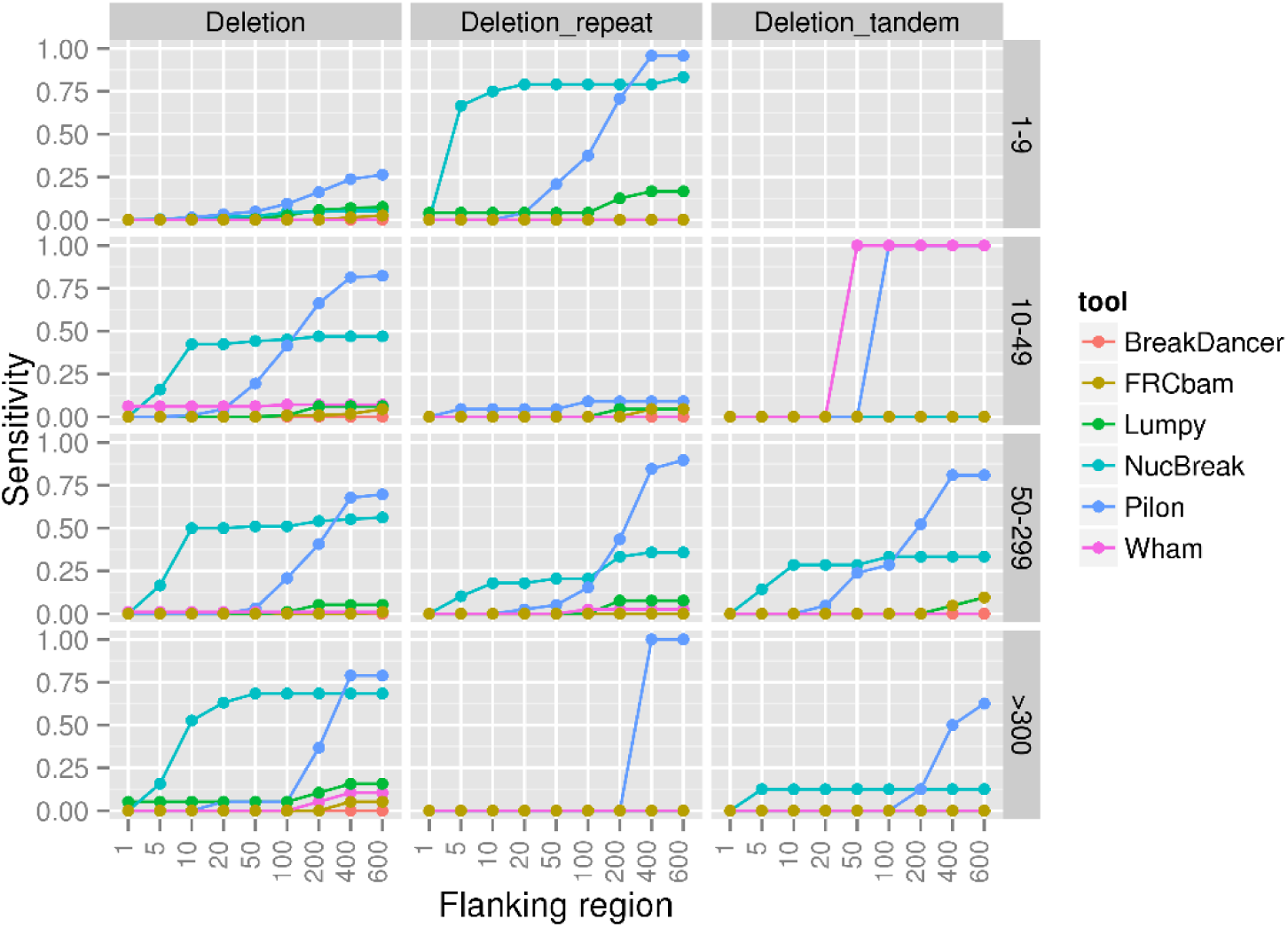
Sensitivity results for the deletion, deletion_repeat and deletion_tandem groups, obtained using the bacterial genome datasets. The deletion_repeat group contains deletions of interspersed repeats or their parts. The deletion_tandem group contains deletions of tandem repeats or their parts.

**Figure 23.**
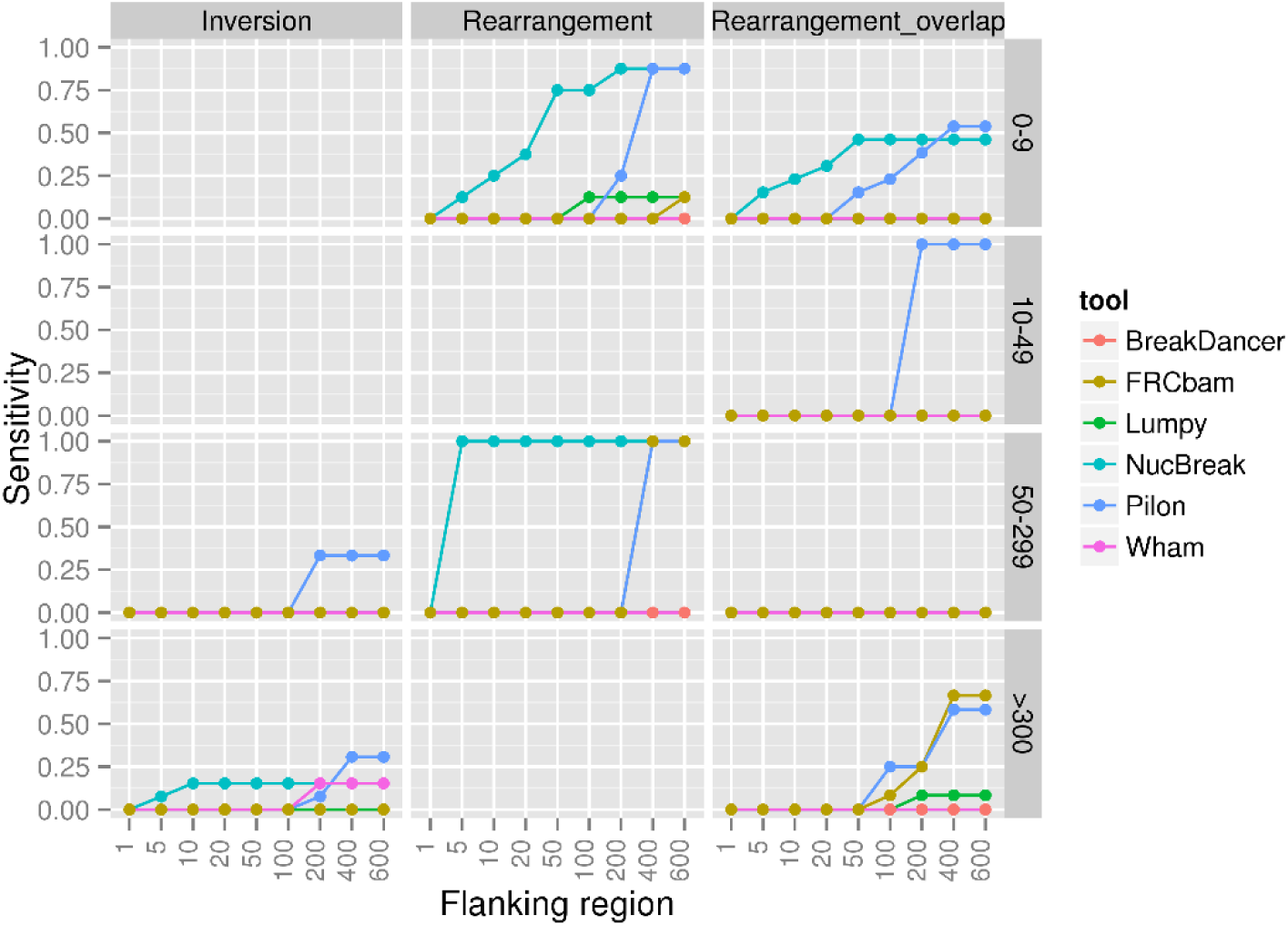
Sensitivity results for the inversion, rearrangement and rearrangement_overlap groups, obtained using the bacterial genome datasets. The rearrangement group consists of relocations and translocations with either inserted regions between misjoined regions (size varied between 1 and 1000) or without them (size is equal to 0). The rearrangement_overlap group consists of relocations and translocations with overlapped misjoined regions.

**Figure 24.**
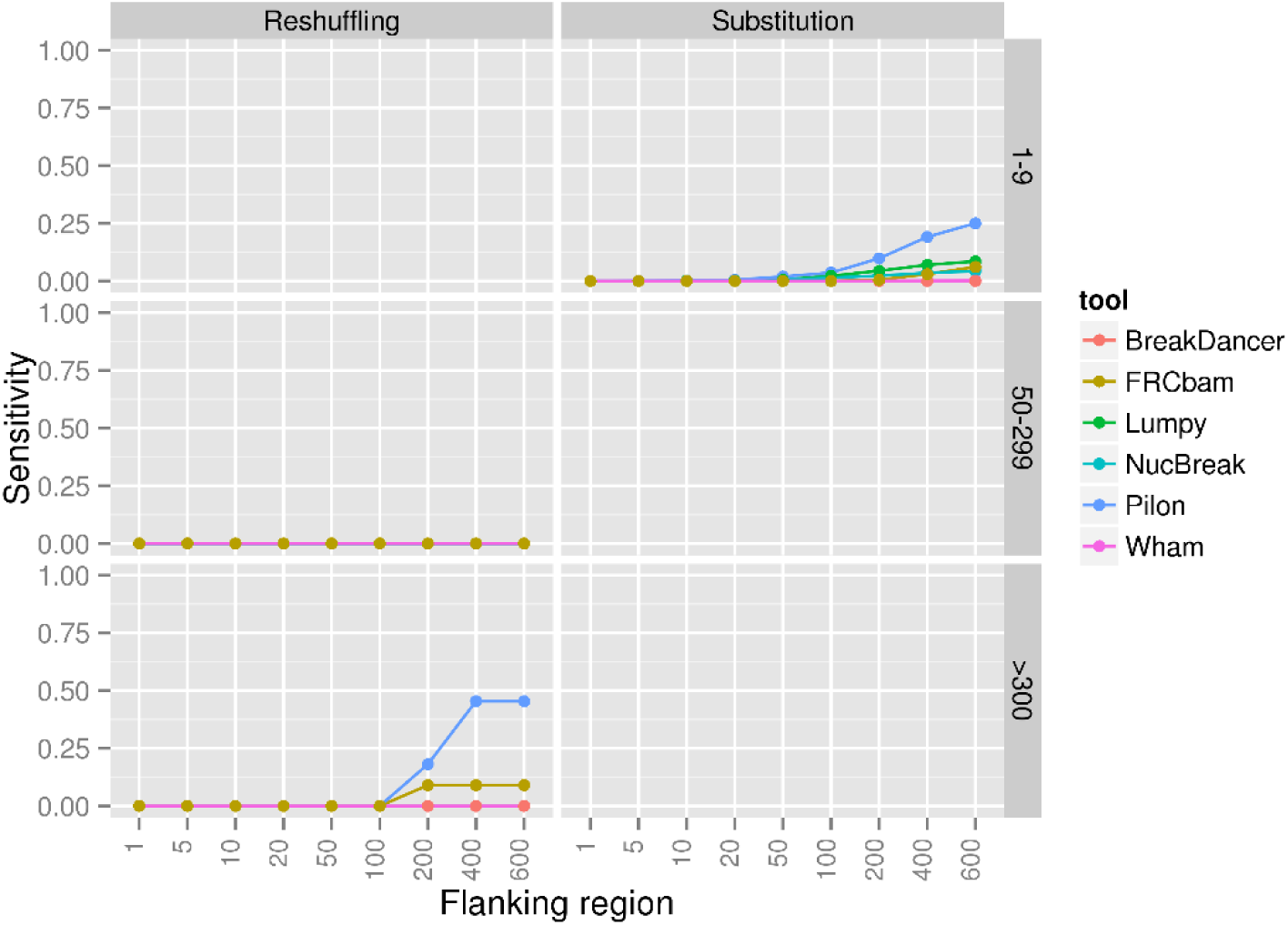
Sensitivity results for the reshuffling and substitution groups, obtained using the bacterial genome datasets.

**Figure 25.**
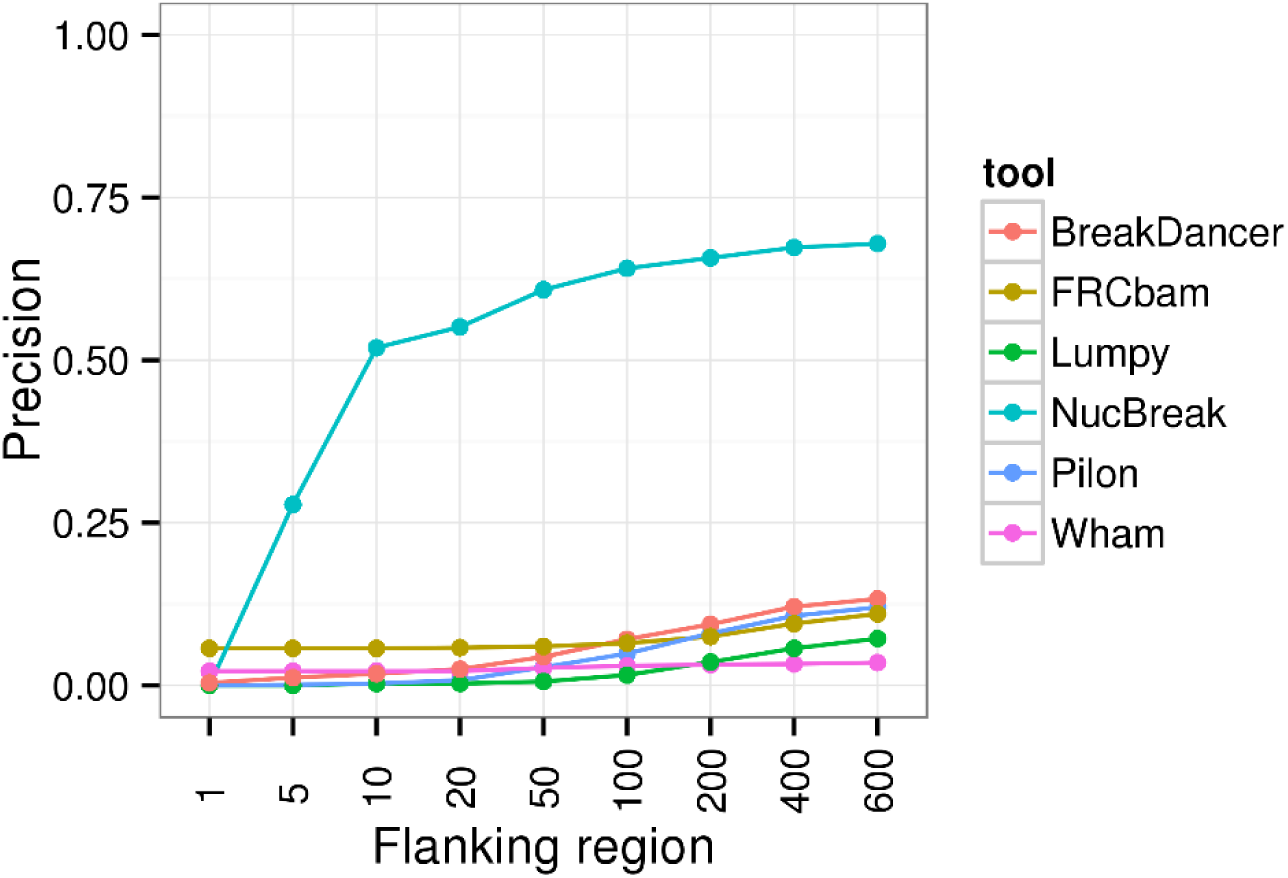
Overall precision obtained using the bacterial genome datasets.

## 4. Discussion

In this paper, we have introduced a tool called NucBreak that detects errors in assemblies by using short paired-end Illumina reads. Neither a reference genome nor a long jump library are required. NucBreak enables detection of assembly errors of all types and sizes, except (1) small insertions, deletions and substitutions that do not change repeat copy numbers, (2) deletions of copies of long interspersed repeats together with bases between repeat copies or long tandem repeat units, and (3) relocations and translocations with long overlapped misjoined regions. The inability of NucBreak to detect such types of assembly errors can be explained by two facts. First, NucBreak does not analyse small differences (approximately up to 30 bp) that are detected during the mapping process, and, thus, misses small insertion, deletion and substitution assembly errors. Second, NucBreak cannot detect errors in the regions that are covered with overlapping properly mapped reads, and, as the result, deletions of copies of long repeats together with bases between repeat copies and rearrangements with long overlapped misjoined regions remain unnoticed. The benchmarking results have shown that NucBreak detects all other assembly errors with high precision and relatively high sensitivity. Such a balance between sensitivity and precision makes NucBreak a good alternative to the existing assembly accuracy assessment tools and SV detection tools.

We have compared NucDiff with several existing tools for assembly accuracy assessment, namely Pilon, FRCbam and REAPR, as well as with some SV detection tools, including BreakDancer, Lumpy and Wham. Only Pilon, REAPR and Wham detect assembly errors of most types and sizes with high sensitivity. However, the high sensitivity of these tools is always combined with low precision. All other tools demonstrate quite low sensitivity and precision, showing good sensitivity results only for some specific assembly error types and sizes.

The results reveal that all tested tools do not output their predictions with a single-nucleotide positional accuracy. All tools obtain better sensitivity when the flanking region size increases. However, Wham and Lumpy do not show such rapid growth of sensitivity as other tools. It means that their initial predictions were more proximal to the annotated assembly errors when at all detected.

It has been also observed that the read coverage is an important factor for detecting structural errors. In the case of REAPR and NucBreak, increase in coverage leads to decrease of sensitivity, while in case of Wham, BreakDancer, and Lumpy it helps to improve sensitivity. The sensitivity of Pilon and FRCBam either decreases or increases with coverage increment, depending on the types and sizes of detected assembly errors.

## 5. Conclusions

We have presented the tool NucBreak aimed at detection of structural errors in assemblies by using Illumina paired-end reads. NucBreak analyses the alignments of reads properly mapped to an assembly and exploits information about alternative read alignments. It enables detection of insertions, deletions, duplications, inversions, and different inter-and intra-chromosomal rearrangements. We have compared NucBreak with REAPR, FRCbam, Pilon, BreakDancer, Lumpy, and Wham. The benchmarking results have shown that in general NucBreak predicts assembly errors with relatively high sensitivity and with higher precision than the other tools. We have also obtained evidence that Lumpy, BreakDancer and Wham, the tools developed for SV detection, can be used for assembly error detection, although in general the sensitivity of these tools, except Wham, is much lower compared to Pilon, REAPR and NucBreak.

## Supporting information

## Abbreviations

SV: structural variant
bp: base pairs

## Declarations

## Acknowledgements

The authors wish to thank the Centre for Ecological and Evolutionary Synthesis (CEES) for access to the computational infrastructure (‘cod’ servers) that enabled the bioinformatics analysis for this project. The authors also wish to thank Karin Lagesen for valuable input in the early phase of the project.

## Availability of data and materials

- Project name: NucBreak
- Project home page: https://github.com/uio-bmi/NucBreak
- Operating system(s): Unix-like system such as Ubuntu Linux and MacOS X.
- Programming language: Python
- Other requirements: Python 2.7
- License: Mozilla Public License (MPL), version 2.0
- Any restrictions to use by non-academics: No
- Additional data: All data used is available as described in Section 2.6

## Authors’ contributions

KK designed and implemented NucBreak. KK, GKS, AJN and TR suggested the demonstration examples and other experiments performed. KK performed all the experiments. KK and TR wrote the manuscript. GKS and AJN revised the manuscript. All authors read and approved the final manuscript.

## Ethics approval and consent to participate

Not applicable.

## Consent for publication

Not applicable.

## Additional files

**Additional file 1** Supplementary materials (PDF 1.02 Mb)

Figure S1 Possible type order and locations of read paths in the absence of breakpoints

Table S1 Genome modifications implemented during the simulation process. G and A denote a reference genome and assembly, respectively. All other letters denote reference genome and assembly sequence regions. Diff means difference. C’ is the reverse complement of C.

Table S2 List of bacterial genomes

Table S3 Number of ground truth errors in each group

