## Supplementary material for "NucBreak: Location of structural errors in a genome assembly by using paired-end Illumina reads"

*Supplementary materials*

Ksenia Khelik, Geir Kjetil Sandve, Alexander Johan Nederbragt, Torbjørn Rognes

### Supplementary methods and results

#### 1. Fragment size estimation

Only read pairs satisfying the following conditions are used for fragment size estimation:

1. Each read in a pair is uniquely aligned
2. Both reads are mapped to the same genome sequence
3. The reads have different orientations relative to the genome sequence
4. The read with the reverse orientation is located at the same position or further down on the sequence compared to the mapping locations of the forward-oriented read
5. The forward- and reverse-oriented reads are not soft-clipped at both sides. However, the alignments of properly mapped reads may contain short substitutions, insertions and deletions.

The fragment size is calculated by the formula:

$frag\_size = ref\_end_2 - ref\_st_1 + 1$ , where

$ref\_end_2$ - the location of reverse-oriented read end at the genome chromosome
$ref\_st_1$ - the location of forward-oriented read start at the genome chromosome

The fragment sizes are sorted in ascending order, and for each fragment size the number of read pairs ($\#P$) having the given fragment size is calculated. Then $min\_frag\_size$ and $max\_frag\_size$ are found:

$$min\_frag\_size = \{frag\_size_i: \#P(frag\_size_i) \geq 10 \; and \; \forall j \; < i \;\; \#P(frag\_size_j) < 10 \\ and \; \exists k = i+1 \ldots i+10 \;\; \#P(frag\_size_k) \geq 10 \; and \; \#k \geq 3\}$$

$$max\_frag\_size = \{frag\_size_i: \#P(frag\_size_i) \geq 10 \; and \; \forall j \; > i \;\; \#P(frag\_size_j) < 10 \\ and \; \exists k = i-11 \ldots i-1 \;\; \#P(frag\_size_k) \geq 10 \; and \; \#k \geq 3\}$$

If the number of corresponding read pairs is less than 10 for any fragment size, then:

$$min\_frag\_size = \{\max(0, frag\_size_i - 50): \forall k \neq i \; \#P(frag\_size_i) \geq \; \#P(frag\_size_k)\}$$
$$max\_frag\_size = \{frag\_size_i \; + 50: \forall k \neq i \; \#P(frag\_size_i) \geq \; \#P(frag\_size_k)\}$$

### 2. Fragment size detection between properly mapped read pairs

Since the reads from properly mapped reads pairs may be soft-clipped in the start or at the end of the read depending on the read orientation, a fragment size inside properly mapped reads is calculated by the extended formula:

$frag\_size = ref\_end_2 + end\_clipped\_dist_2 - ref\_st_1 - start\_clipped\_dist_1 + 1$, where

$ref\_end_2$ - the location of the reverse-oriented read end at the genome chromosome
$end\_clipped\_dist_2$ - the number of soft-clipped bases at the end of the reverse-oriented read
$ref\_st_1$ - the location of the forward-oriented read start at the genome chromosome
$start\_clipped\_dist_1$- the number of soft-clipped bases in the beginning of the reverse-oriented read

### 3. The Velvet, Abyss and Spades parameter settings used to obtain assemblies

Spades was run with the “-t 2 -k 33 --cov-cutoff 2” parameter settings.
Abyss was run with “k=64” parameter setting.
Velveth was run with k-mer length equal to 31.
Velvetg was run with “-ins_length 180 -scaffolding yes -min_contig_lgth 250 -cov_cutoff 5” parameter settings.

### 4. The NucBreak, REAPR and FRCbam parameter settings used to detect assembly errors

In the Sections 3.1 and 3.2, we used the following parameter settings for the tools:

- NucBreak was run with “--min_frag_size 620 --max_frag_size 790” parameter settings
- In case of REAPR, perfectmap was run with 700 bp average insert size
- FRCbam was run with “--pe-max-insert 776” and the value for “--genome-size” parameter was detected automatically by using python script for each modification case.

In the Section 3.3, we used the following parameter settings for the tools:

- in case of REAPR, perfectmap was run with 300 bp average insert size
- FRCbam was run with "--pe-max-insert 776 --genome-size 112000000" parameter settings

In the Section 3.4, we used the following parameter settings for the tools:

- In case of REAPR, perfectmap was run with the following average insert sizes depending on the genome dataset used:
    - Salmonella dataset - 500 bp
    - Staphylococcus dataset - 400 bp
    - Escherichia dataset - 300 bp
    - Pseudomonas dataset - 180 bp
    - Bordetella dataset - 450 bp
    - Brucella dataset - 500 bp
    - Klebsiella dataset - 200 bp
    - Enterobacter dataset - 300 bp
- FRCbam was run with the following parameter settings depending on the genome dataset used:
    - Salmonella dataset - "--pe-max-insert 1060 --genome-size 4810000"
    - Staphylococcus dataset - "--pe-max-insert 1040 --genome-size 2860000"
    - Escherichia dataset - "--pe-max-insert 1110 --genome-size 5480000"
    - Pseudomonas dataset - "--pe-max-insert 844 --genome-size 6820000"
    - Bordetella dataset - "--pe-max-insert 890 --genome-size 4110000"
    - Brucella dataset - "--pe-max-insert 1120 --genome-size 3300000"
    - Klebsiella dataset - "--pe-max-insert 950 --genome-size 5720000"
    - Enterobacter dataset - "--pe-max-insert 819 --genome-size 5040000"

### 5. Result evaluation

The ground truth entries may be represented as dots (e.g. in case of deletions, simple relocations or translocations) or as intervals (e.g. in case of insertion, duplications, relocations with overlap). If a ground truth entry is an interval, it may be fully covered with reads mapped back to the query sequences (e.g. in case of inversions) or remain uncovered (e.g. in case of inserted regions that are not present in the reference genome). In the first case, a tested tool is expected to mark the regions corresponding to the start- and end-points of the ground truth entry as breakpoints, while in the second case the whole entry is expected to be predicted as a breakpoint.

We say that if a ground truth entry coincides with an obtained breakpoint or the ground truth entry start- and end-points both coincide with obtained breakpoints, then we have a true positive (TP). If a ground truth entry does not coincide with any of obtained breakpoints, or either the start- or end-point coincides with an obtained breakpoint, then we have a false negative (FN). To get TPs and FNs, we have run BEDTools with the pairtopair -both' option. With this option, BEDTool reports an overlap between two intervals A and B if both ends of A overlap B. If BEDTool reports an overlap for a whole ground truth entry or for its start- and end-points, then we get a TP, otherwise a FN. Having obtained the number of TPs and FNs, we calculate sensitivity by the formula:

$$Sensitivity = \frac{\#TP}{\#TP + \#FN}$$

Unlike ground truth entries, an obtained result can correspond only to one interval: either to a whole ground truth entry or to its start- or end-point. We say that if an obtained breakpoint coincides with a whole ground truth entry or with either its start- or end-point, then the given obtained breakpoint is a true positive (TP*). Since two true positive obtained breakpoints may correspond to one true positive ground truth entry (in the case when both the start- and end-point of a ground truth entry is correctly predicted by a tool), the numbers of TPs and TP*s are not equal, and the number of TPs cannot be used to calculate precision. If an obtained breakpoint does not coincide with any of the ground truth entries and with any of the ground truth entry start- and end-points, then the given obtained breakpoint is a false positive (FP). To get TP* and FP, we have run BEDTools with the ‘‘pairtopair -notboth’ option. With this option, BEDTool reports an overlap between two intervals A and B, if one or neither of A's ends overlap B. If BEDTool reports an overlap for an obtained breakpoint with a whole ground truth entry or with its ends, then we get a FP, otherwise a TP*. Having obtained the number of TP*s and FPs, we calculate precision by the formula:

$$Precision = \frac{\#TP^*}{\#TP^* + \#FP}$$

### Supplementary figures

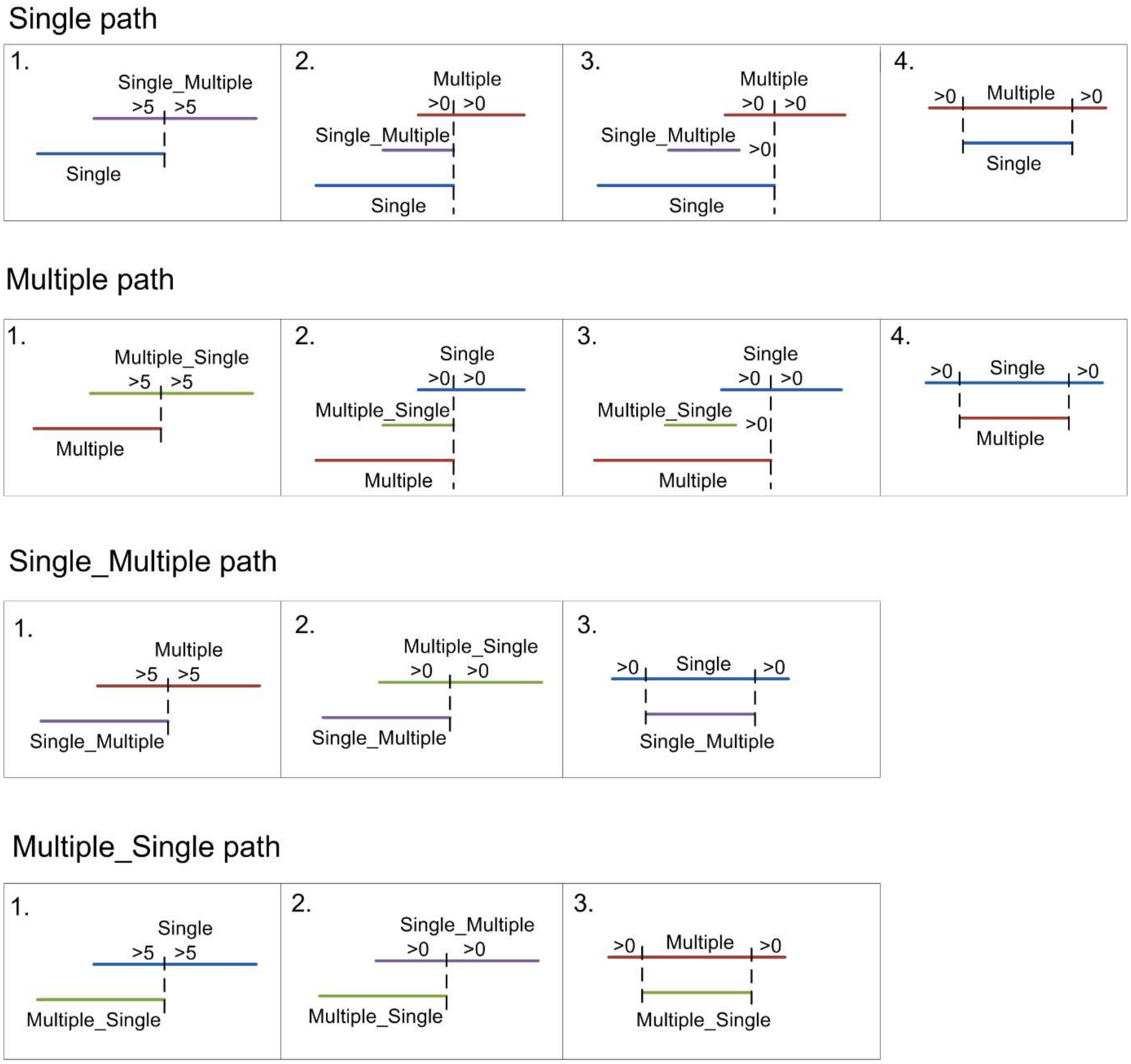


**Figure S1** Possible type order and locations of read paths in the absence of breakpoints.

### Supplementary tables

**Table S1** Genome modifications implemented during the simulation process. G and A denote a reference genome and assembly, respectively. All other letters denote reference genome and assembly sequence regions. Diff means difference. C' is the reverse complement of C.

| **Insertions** | |
|---|---|
| 1. G: HB<br>A: HCB<br>Diff: insertion | 46. G: DLLLLxC<br>A: DCLLLLxC<br>Diff: duplication |
| 2. G: HBxC<br>A: HCBxC<br>Diff: duplication | 47. G: DLLLLxTKxTKxTK<br>A: DTLLLLxTKxTKxTK<br>Diff: duplication |
| 3. G: HBxTKxTKxTK<br>A: HTBxTKxTKxTK<br>Diff: duplication | 48. G: DLLLLxTKxTKxTK<br>A: DKLLLLxTKxTKxTK<br>Diff: duplication |
| 4. G: HBxCxCxC<br>A: HCBxCxCxC<br>Diff: duplication | 49. G: DLLLLxCxCxC<br>A: DCLLLLxCxCxC<br>Diff: duplication |
| 5. G: HBxTKTKTKTK<br>A: HTBxTKTKTKTK<br>Diff: duplication | 50. G: DKTKTKTKT<br>A: DTKTKTKTKT<br>Diff: duplication |
| 6. G: HBxCCCC<br>A: HCBxCCCC<br>Diff: duplication | 51. G: DKTKTKTKT<br>A: DKKTKTKTKT<br>Diff: tandem_duplication |
| 7. G: HBxTKTKTKTK<br>A: HKTBxTKTKTKTK<br>Diff: duplication | 52. G: DCCCC<br>A: DCCCCC<br>Diff: tandem_duplication |
| 8. G: HBxCCCC<br>A: HCCCCBxCCCC<br>Diff: duplication | 53. G: DKTKTKTKT<br>A: DTKKTKTKTKT<br>Diff: duplication |
| 9. G: RxRxR<br>A: RxCRxR<br>Diff: insertion | 54. G: DPPPPxTKTKTKTK<br>A: DTPPPPxTKTKTKTK<br>Diff: duplication |
| 10. G: RxRxRxC<br>A: RxCRxRxC<br>Diff: duplication | 55. G: DPPPPxTKTKTKTK<br>A: DKPPPPxTKTKTKTK<br>Diff: duplication |
| 11. G: TKxTKxTK<br>A: TKxTTKxTK<br>Diff: tandem_duplication | 56. G: DPPPPxCCCC<br>A: DCPPPPxCCCC<br>Diff: duplication |
| 12. G: TKxTKxTK<br>A: TKxKTKxTK<br>Diff: duplication | 57. G: DPPPPxTKTKTKTK<br>A: DKTPPPPxTKTKTKTK<br>Diff: duplication |

13. G: CxCxC
A: CxCCxC
Diff: tandem_duplication

14. G: RxRxRxTKTKTKTK
A: RxTRxRxTKTKTKTK
Diff: duplication

15. G: RxRxRxTKTKTKTK
A: RxKRxRxTKTKTKTK
Diff: duplication

16. G: RxRxRxTKTKTKTK
A: RxTKRxRxTKTKTKTK
Diff: duplication

17. G: RxRxRxTKTKTKTK
A: RxKTRxRxTKTKTKTK
Diff: duplication

18. G: RxRxRxCCCC
A: RxCCCCRxRxCCCC
Diff: duplication

19. G: RxRxRxTKxTKxTK
A: RxTRxRxTKxTKxTK
Diff: duplication

20. G: RxRxRxTKxTKxTK
A: RxKRxRxTKxTKxTK
Diff: duplication

21. G: RxRxRxCxCxC
A: RxCRxRxCxCxC
Diff: duplication

22. G: RxRxR
A: RxRCxR
Diff: insertion

23. G: RxRxRxC
A: RxRCxRxC
Diff: duplication

24. G: TKxTKxTK
A: TKxTKTxTK
Diff: duplication

25. G: TKxTKxTK
A: TKxTKKxTK
Diff: tandem_duplication

26. G: RxRxRxTKTKTKTK
A: RxRTxRxTKTKTKTK
Diff: duplication

27. G: RxRxRxTKTKTKTK
A: RxRKxRxTKTKTKTK
Diff: duplication

58. G: DPPPPxCCCC
A: DCCCCPPPPxCCCC
Diff: duplication

59. G: LLLLD
A: LLLLCD
Diff: insertion

60. G: LLLLDxC
A: LLLLCDxC
Diff: duplication

61. G: LLLLDxTKxTKxTK
A: LLLLTDxTKxTKxTK
Diff: duplication

62. G: LLLLDxTKxTKxTK
A: LLLLKDxTKxTKxTK
Diff: duplication

63. G: LLLLDxCxCxC
A: LLLLCDxCxCxC
Diff: duplication

64. G: TKTKTKTKD
A: TKTKTKTKTD
Diff: duplication

65. G: TKTKTKTKD
A: TKTKTKTKKD
Diff: tandem_duplication

66. G: TKTKTKTKD
A: TKTKTKTKKTD
Diff: duplication

67. G: PPPPDxTKTKTKTK
A: PPPPTDxTKTKTKTK
Diff: duplication

68. G: PPPPDxTKTKTKTK
A: PPPPKDxTKTKTKTK
Diff: duplication

69. G: PPPPDxCCCC
A: PPPPCDxCCCC
Diff: duplication

70. G: PPPPDxTKTKTKTK
A: PPPPKTDxTKTKTKTK
Diff: duplication

71. G: PPPPDxCCCC
A: PPPPCCCCDxCCCC
Diff: duplication

72. G: PPPP
A: PPCPP
Diff: insertion

| | |
|---|---|
| 28. G: RxRxRxTKTKTKTK<br>A: RxRTKxRxRxTKTKTKTK<br>Diff: duplication | 73. G: PPPPxC<br>A: PPCPPxC<br>Diff: duplication |
| 29. G: RxRxRxTKTKTKTK<br>A: RxRDTxRxRxTKTKTKTK<br>Diff: duplication | 74. G: PPPPxTKxTKxTK<br>A: PPTPPxTKxTKxTK<br>Diff: duplication |
| 30. G: RxRxRxCCCC<br>A: RxRCCCCxRxCCCC<br>Diff: duplication | 75. G: PPPPxTKxTKxTK<br>A: PPKPPxTKxTKxTK<br>Diff: duplication |
| 31. G: RxRxRxTKxTKxTK<br>A: RxRTxRxTKxTKxTK<br>Diff: duplication | 76. G: PPPPxCxCxC<br>A: PPCPPxCxCxC<br>Diff: duplication |
| 32. G: RxRxRxTKxTKxTK<br>A: RxRKxRxTKxTKxTK<br>Diff: duplication | 77. G: TKTKTKTK<br>A: TKTKKTKTK<br>Diff: tandem_duplication |
| 33. G: RxRxRxCxCxC<br>A: RxRCxRxCxCxC<br>Diff: duplication | 78. G: PPPPxTKTKTK<br>A: PPTPPxTKTKTK<br>Diff: duplication |
| 34. G: RDxRDxRD<br>A: RDxRCDxRD<br>Diff: insertion | 79. G: PPPPxTKTKTK<br>A: PPKPPxTKTKTK<br>Diff: duplication |
| 35. G: RDxRDxRDxC<br>A: RDxRCDxRDxC<br>Diff: duplication | 80. G: PPPPxCCC<br>A: PPCPPxCCC<br>Diff: duplication |
| 36. G: TKxTKxTK<br>A: TKxTKKxTK<br>Diff: tandem_duplication | 81. G: PPPPxTKTKTK<br>A: PPKTPPxTKTKTK<br>Diff: duplication |
| 37. G: RDxRDxRDxTKTKTKTK<br>A: RDxRTDxRDxTKTKTKTK<br>Diff: duplication | 82. G: PPPPxCCC<br>A: PPCCCPPxCCC<br>Diff: duplication |
| 38. G: RDxRDxRDxTKTKTKTK<br>A: RDxRKDxRDxTKTKTKTK<br>Diff: duplication | 83. G: DKL<br>A: DKTKL<br>Diff: insertion |
| 39. G: RDxRDxRDxCCCC<br>A: RDxRCDxRDxCCCC<br>Diff: duplication | 84. G: DKLxT<br>A: DKTKLxT<br>Diff: insertion |
| 40. G: RDxRDxRDxTKTKTKTK<br>A: RDxRKTDxRDxTKTKTKTK<br>Diff: duplication | 85. G: LxKLxLxT<br>A: LxKTKLxLxT<br>Diff: insertion |
| 41. G: RDxRDxRDxCCCC<br>A: RDxRCCCCDxRDxCCCC<br>Diff: duplication | 86. G: KLxKLxKLxT<br>A: KLxKTKLxKLxT<br>Diff: insertion |
| 42. G: RDxRDxRDxTKxTKxTK<br>A: RDxRTDxRDxTKxTKxTK<br>Diff: duplication | 87. G: LKxLKxLKxT<br>A: LKxLKTKxLKxT<br>Diff: insertion |

| | |
|---|---|
| 43. G: RDxRDxRDxTKxTKxTK<br>A: RDxRKDxRDxTKxTKxTK<br>Diff: duplication | 88. G: DCR<br>A: DCCR<br>Diff: tandem_duplication |
| 44. G: RDxRDxRDxCxCxC<br>A: RDxRCDxRDxCxCxC<br>Diff: duplication | 89. G: DCR<br>A: DCCCCR<br>Diff: tandem_duplication |
| 45. G: DLLLL<br>A: DCLLLL<br>Diff: insertion | 90. G: LxCLxL<br>A: LxCCLxL<br>Diff: tandem_duplication |
| **Deletions** | |
| 1. G: RCD<br>A: RD<br>Diff: deletion | 14. G: DTKKKK<br>A: DKKKK<br>Diff: deletion |
| 2. G: RxCRxR<br>A: RxRxR<br>Diff: deletion | 15. G: DTKKKK<br>A: DKKK<br>Diff: deletion |
| 3. G: KRxTKRxKR<br>A: KRxRxKR<br>Diff: deletion_repeat | 16. G: DTKKKK<br>A: DK<br>Diff: deletion_tandem |
| 4. G: RxCRxR<br>A: RxxR<br>Diff: deletion_repeat | 17. G: DTKKKKF<br>A: DF<br>Diff: deletion |
| 5. G: KxTKFxK<br>A: KxxK<br>Diff: deletion_repeat | 18. G: DTLLLLK<br>A: D<br>Diff: deletion |
| 6. G: CRxCRxCR<br>A: CRxRxCR<br>Diff: deletion_repeat | 19. G: RTKLKLKLKL<br>A: RLKLKLKL<br>Diff: deletion |
| 7. G: CxCxC<br>A: CxxC<br>Diff: deletion_repeat | 20. G: DTKTKTKTK<br>A: DKTKTKTK<br>Diff: deletion |
| 8. G: KxKTxK<br>A: KxxK<br>Diff: deletion_repeat | 21. G: DTKTKTKTK<br>A: DTKTTKTK<br>Diff: deletion |
| 9. G: KTxKTxKT<br>A: KTxKxKT<br>Diff: deletion_repeat | 22. G: TKTKTKTKD<br>A: TKTKTKTD<br>Diff: deletion |
| 10. G: RTxRTKxRT<br>A: RTxRxRT<br>Diff: deletion_repeat | 23. G: DCCCC<br>A: DCCC<br>Diff: deletion_tandem |
| 11. G: RxRCxR<br>A: RxRxR<br>Diff: deletion | 24. G: RCCCCD<br>A: RD<br>Diff: deletion |
| 12. G: KTKxK<br>A: KxK<br>Diff: deletion_repeat | 25. G: DTTTTK<br>A: DTTT<br>Diff: deletion |

| | |
|---|---|
| 13. G: KTK<br>A: K<br>Diff: deletion_repeat | 26. G: DTTTTK<br>A: D<br>Diff: deletion |
| **Relocations** | **Inversions** |
| 1. G: SzV<br>A: SCV<br>Diff: relocation<br><br>2. G: SzV<br>A: SV<br>Diff: relocation<br><br>3. G: SCzCV<br>A: SCV<br>Diff: relocation_overlap | 1. G: DCR<br>A: DC'R<br>Diff: inversion<br><br>2. G: DCRxRxR<br>A: DC'RxRxR<br>Diff: inversion<br><br>3. G: DRCxRxR<br>A: DRC'xRxR<br>Diff: inversion<br><br>4. G: RCDxRCDxRCD<br>A: RCDxRC'DxRCD<br>Diff: inversion |

In the simulated modifications, the following lengths of regions were used:

1. Distance between each manipulation case =2500 bp
2. len(H)=800 bp
3. len(B)=800 bp
4. len(x)=800 bp
5. len(C)={17,30,100,250,800} bp
6. len(TK)={[50,70],[100,150],[250,250],[600,600]} bp, where first number in a pair is len(T) and second number in a pair is len(K)
7. len(R)=600 bp
8. len(D)=600 bp
9. len(L)=200 bp
10. len(P)=400 bp
11. len(F)=len(T) in len(TK)
12. len(S)=1500 bp
13. len(V)=1500 bp
14. len(Z)=15000 bp

**Table S2** List of bacterial genomes.

| Genome | Genome length, Mb | Accession number | Reads length, bp (first, second) | Coverage | Read library accession number |
|---|---|---|---|---|---|
| *Bordetella pertussis* str. J081 | 4,11 | GCA_002859625.1 | 250<br>250 | 32x | SRR5829829 |
| *Brucella melitensis* str. 1 | 3,30 | GCA_900236405.1 | 243 ± 28.8<br>243 ± 28.7 | 40x | ERR2192800 |
| *Enterobacter cloacae* str. AR_0136 | 5,04 | GCA_002204775.1 | 233 ± 34.9<br>233 ± 34.8 | 23x | SRR4025988 |
| *Escherichia coli* str. 2014C-3599 | 5,48 | GCA_003018935.1 | 236 ± 39.0<br>236 ± 38.8 | 60x | SRR1609862 |
| *Klebsiella pneumonia* str. SGH10 | 5,72 | GCA_002813595.1 | 146 ± 15.8<br>146 ± 15.7 | 32x | SRR5082357 |
| *Pseudomonas aeruginosa* str. AR_0095 | 6.82 | GCA_002997005.1 | 229 ± 38.2<br>229 ± 36.9 | 60x | SRR3242025 |
| *Salmonella enterica* str. CFSAN047866 | 4,81 | GCA_003073535.1 | 244 ± 27.3<br>244 ± 27.3 | 37x | SRR3272258 |
| *Staphylococcus aureus* str. CFSAN007896 | 2,86 | GCA_003031425.1 | 236 ± 41.8<br>236 ± 41.7 | 28x | SRR5912676 |

**Table S3** Number of ground truth errors in each group.

| Error type | Error size | Simulated datasets | Assemblathon 1 dataset | Bacterial genome datasets |
|---|---|---|---|---|
| insertion | 0-9 | 0 | 1663 | 402 |
| | 10-49 | 126 | 892 | 414 |
| | 50-299 | 216 | 38 | 133 |
| | >300 | 153 | 31 | 15 |
| duplication | 0-9 | 0 | 23 | 4 |
| | 10-49 | 342 | 1 | 12 |
| | 50-299 | 1359 | 11 | 5 |
| | >300 | 756 | 287 | 1 |

| | | | | |
|---|---|---|---|---|
| tandem_duplication | 0-9 | 0 | 0 | 0 |
| | 10-49 | 54 | 23 | 3 |
| | 50-299 | 234 | 93 | 14 |
| | >300 | 153 | 683 | 0 |
| deletion | 0-9 | 0 | 1833 | 437 |
| | 10-49 | 54 | 1091 | 113 |
| | 50-299 | 243 | 307 | 96 |
| | >300 | 243 | 527 | 19 |
| deletion_repeat | 0-9 | 0 | 424 | 24 |
| | 10-49 | 36 | 10 | 22 |
| | 50-299 | 144 | 21 | 39 |
| | >300 | 207 | 7 | 2 |
| deletion_tandem | 0-9 | 0 | 1 | 0 |
| | 10-49 | 18 | 37 | 1 |
| | 50-299 | 27 | 147 | 21 |
| | >300 | 36 | 24 | 8 |
| inversion | 0-9 | 0 | 2 | 0 |
| | 10-49 | 72 | 0 | 0 |
| | 50-299 | 72 | 6 | 3 |
| | >300 | 36 | 94 | 13 |
| relocation/ rearrangement | 0-9 | 45 | 749 | 8 |
| | 10-49 | 18 | 2 | 0 |
| | 50-299 | 18 | 17 | 1 |
| | >300 | 9 | 95 | 0 |
| relocation_overlap/ rearrangement_overlap | 0-9 | 0 | 744 | 13 |
| | 10-49 | 18 | 25 | 1 |
| | 50-299 | 18 | 152 | 1 |
| | >300 | 9 | 76 | 12 |
| reshuffling | 0-9 | 0 | 4 | 0 |
| | 10-49 | 0 | 1 | 0 |
| | 50-299 | 0 | 11 | 1 |

| | | | | |
|---|---|---|---|---|
| | >300 | 0 | 94 | 11 |
| substitution | 0-9 | 0 | 29 | 8000 |
| | 10-49 | 0 | 1 | 0 |
| | 50-299 | 0 | 3 | 0 |
| | >300 | 0 | 21 | 0 |
